# Purkinje-Enriched snRNA-seq in SCA7 Cerebellum Reveals Zebrin Identity Loss as a Central Feature of Polyglutamine Ataxias

**DOI:** 10.1101/2023.03.19.533345

**Authors:** Luke C. Bartelt, Pawel M. Switonski, Grażyna Adamek, Juliana Carvalho, Lisa A. Duvick, Sabrina I. Jarrah, Hayley S. McLoughlin, Daniel R. Scoles, Stefan M. Pulst, Harry T. Orr, Court Hull, Craig B. Lowe, Albert R. La Spada

## Abstract

Spinocerebellar ataxia type 7 (SCA7) is an inherited neurodegenerative disorder caused by a CAG-polyglutamine repeat expansion. SCA7 patients display a striking loss of Purkinje cell (PC) neurons with disease progression; however, PCs are rare, making them difficult to characterize. We developed a PC nuclei enrichment protocol and applied it to single-nucleus RNA-seq of a SCA7 knock-in mouse model. Our results unify prior observations into a central mechanism of cell identity loss, impacting both glia and PCs, driving accumulation of inhibitory synapses and altered PC spiking. Zebrin-II subtype dysregulation is the predominant signal in PCs, leading to complete loss of zebrin-II striping at motor symptom onset in SCA7 mice. We show this zebrin-II subtype degradation is shared across Polyglutamine Ataxia mouse models and SCA7 patients. It has been speculated that PC subtype organization is critical for cerebellar function, and our results suggest that a breakdown of zebrin-II parasagittal striping is pathological.

## INTRODUCTION

CAG-polyglutamine (polyQ) repeat diseases are a group of nine genetic neurodegenerative disorders caused by the expansion of a CAG trinucleotide repeat in the coding exon of different genes^1^. The CAG-polyQ repeat diseases share common features of protein aggregation and selective neuronal vulnerability, but differ in their clinical manifestations and affected brain regions. These diseases are an important category of neurodegenerative diseases for which highly representative mouse models have been generated and characterized^2–6^, which provide opportunities to discover molecular mechanisms of CAG-polyQ neurodegeneration while also providing valuable insight to sporadic neurodegenerative disorders. Spinocerebellar ataxia type 7 (SCA7) is one CAG-polyQ repeat expansion disease which has been successfully modeled in mice by knock-in of an expanded 266 CAG repeat into the endogenous mouse ataxin-7 locus. SCA7 266Q knock-in mice develop early onset, rapidly progressive cerebellar and retinal degeneration^2^, akin to juvenile-onset SCA7 in human patients^7–9^.

While mouse models have implicated a lengthy list of biological pathways in the SCAs, transcription dysregulation has emerged as a predominant pathogenic theme across the CAG-polyQ repeat diseases, such that these disorders are categorized as “transcriptionopathies”^10^. This shared impact on transcription is typically mediated via the nuclear accumulation of toxic polyQ aggregates, which can sequester important transcription regulatory proteins, such as TATA-binding protein and CREB-binding protein^11,12^. Transcriptional mechanisms are particularly relevant in SCA7, as the ataxin-7 protein performs its primary role in the nucleus as a core component of the STAGA transcription co-activator complex^13,14^, which facilitates RNA Polymerase II-mediated transcription genome-wide^15^. The STAGA complex has several functional domains, including a histone acetyltransferase module that catalyzes the deposition of the activating H3K9ac mark and a deubiquitinase (DUB) module that removes H2BK120ub modifications – tuning both the structural and transcriptional properties of target genes^16–18^. Ataxin-7 sits in the DUB module, and polyQ repeat expansion leads to structural impairments of the STAGA complex and alters global levels of both H3K9ac and H2BK120ub modifications, resulting in widespread transcription dysregulation^19–21^. Interestingly, the transcriptional signatures associated with SCA7 cerebellar degeneration significantly correlate with differential gene expression changes in the cerebellum of SCA1 and SCA2 mouse models, implying the existence of common transcriptional impacts by different polyQ-expanded ataxin proteins, despite non-overlapping normal functions^22–24^. The basis for highly shared patterns of gene expression alteration in the polyglutamine cerebellar degenerations remains unknown.

The cerebellum is a highly organized, yet complex region of the central nervous system (CNS) with over 15 distinct cell types coordinating together to maintain proper cerebellar function^25^. The action of these cells is organized around the cerebellar Purkinje cell (PC), which is the largest and most metabolically active neuron in the CNS^26,27^, and serves as the sole output neuron of the cerebellar cortex. Purkinje cell dysfunction and neurodegeneration are central to the development of ataxic phenotypes in SCA7 mice and human patients^28^. However, numerous studies have also demonstrated the critical involvement of Bergmann glia and cerebellar astrocytes in SCA7 Purkinje cell degeneration, indicating that there is a non-cell autonomous contribution to SCA7 Purkinje cell degeneration^29,30^. Furthermore, Purkinje neurons exist as two sub-types: zebrin-II-positive and zebrin-II-negative PCs, organized along parasagittal stripes in the cerebellum, each with distinct gene expression signatures, synaptic organization, electrophysiology, and circuitry^31–34^. While several molecular and functional features of these distinct zebrin-II zones have been characterized, it remains unclear if these zones are essential for proper cerebellar processing and function.

To determine the relative contribution of cerebellar cell types and subtypes to SCA7 pathogenesis, we pursued a single-nucleus RNA-seq analysis of SCA7 266Q knock-in mice. Although single-cell and single-nucleus sequencing technologies offer the ability to isolate transcriptional signals of diverse cell types in tissues such as the cerebellum, standard methods suffer from unique challenges in the cerebellum due to the massive overabundance of granule cell neurons. To overcome this issue, we developed a simple and robust Purkinje cell nuclear enrichment protocol for use in single-nucleus sequencing assays, yielding an enrichment of Purkinje cells from 1% to >40% of the captured cell population in adult mouse cerebellum. The results of our snRNA-seq analysis of SCA7 266Q knock-in mouse cerebellum revealed significant expression alterations involving synapse organization genes, which we further investigated by directly examining synapse distribution and circuit function in SCA7 mice. In addition, we detected a striking alteration of SCA7 PC zebrin-II subtype specification at the level of gene expression, and upon anatomical analysis, we discovered a complete ablation of zebrin-II parasagittal striping in SCA7 knock-in mice. When we evaluated zebrin-II parasagittal striping patterns in related polyglutamine SCAs, we discovered that this dysregulation of PC zebrin-II subtypes extends to other polyglutamine disease mouse models, suggesting that dysfunctional zebrin-II subtype maintenance is a defining feature of polyglutamine ataxias.

## RESULTS

### Purkinje cell-enriched snRNA-seq yields a high-resolution map of the SCA7 cerebellum

To enrich for PCs, which are central to degeneration of the cerebellum in SCA7 and related ataxias, we developed a nuclear enrichment protocol which utilizes a modified nuclei isolation buffer with increased concentration of K_2_SO_4_ to preferentially increase the proportion of PC nuclei in suspension (**Figure S1**). With this PC nuclear enrichment method, we then pursued a single-nucleus RNA-seq (snRNA-seq) analysis of cerebella from SCA7 266Q knock-in mice and wild-type (WT) littermate controls (**Figure 1A**), at two successive timepoints: 5 weeks of age (presymptomatic) and 8.5 weeks of age (early symptomatic). To minimize batch effects and account for any sex differences, two male and two female mice for each genotype were barcoded and multiplexed with MULTI-seq cholesterol-modified oligonucleotides^35^, and run in a single 10X well. After quality control, we recovered 5334 and 8820 nuclear profiles from the 5-week and 8.5-week timepoints respectively, which were assigned to their animal of origin by MULTI-seq barcode counts. UMAP clustering separated nuclei into known resident cell types of the cerebellum with a significant enrichment of Purkinje cells to >40% of recovered nuclei (**Figure 1B**). Nuclei originating from WT and SCA7 mice displayed high levels of concordance in 5 week-old mice with relatively few Differentially Expressed Genes (DEGs) limited only to PCs (**Supplementary Table 1**). However, at 8.5 weeks of age, we identified dramatic transcriptional differences between WT and SCA7 mice, visually represented by the genotypes separately clustering in UMAP space and quantified by large numbers of DEGs in PCs, oligodendrocytes, Bergmann glia, and astrocytes (**Figure 1C-D; Supplementary Table 2**). Immediately apparent in the differential expression was a bias toward downregulation of genes across SCA7 cell types (**Figure 1E-F**). Furthermore, due to our controlled multiplexed experimental design, we were able to assess differential RNA abundance between SCA7 and WT animals. We observed a cell-type selective decrease in nuclear RNA abundance in affected cell-types, most apparent in PCs at both the 5-week and 8.5-week timepoints (**Figure S3**). Given this significant difference in RNA abundance and the known role of the STAGA complex in transcription initiation genome-wide, proper differential expression testing is warranted for analysis of this single-nucleus dataset, as standard log-normalization of gene counts may yield misleading results in the context of global transcriptional effects^36^. Therefore, we performed a replicate structured pseudobulk analysis of gene counts between SCA7 and WT cell types with DEseq2, removing the standard assumption that total RNA abundance stays constant across groups being compared^37,38^.

**Figure 1:**
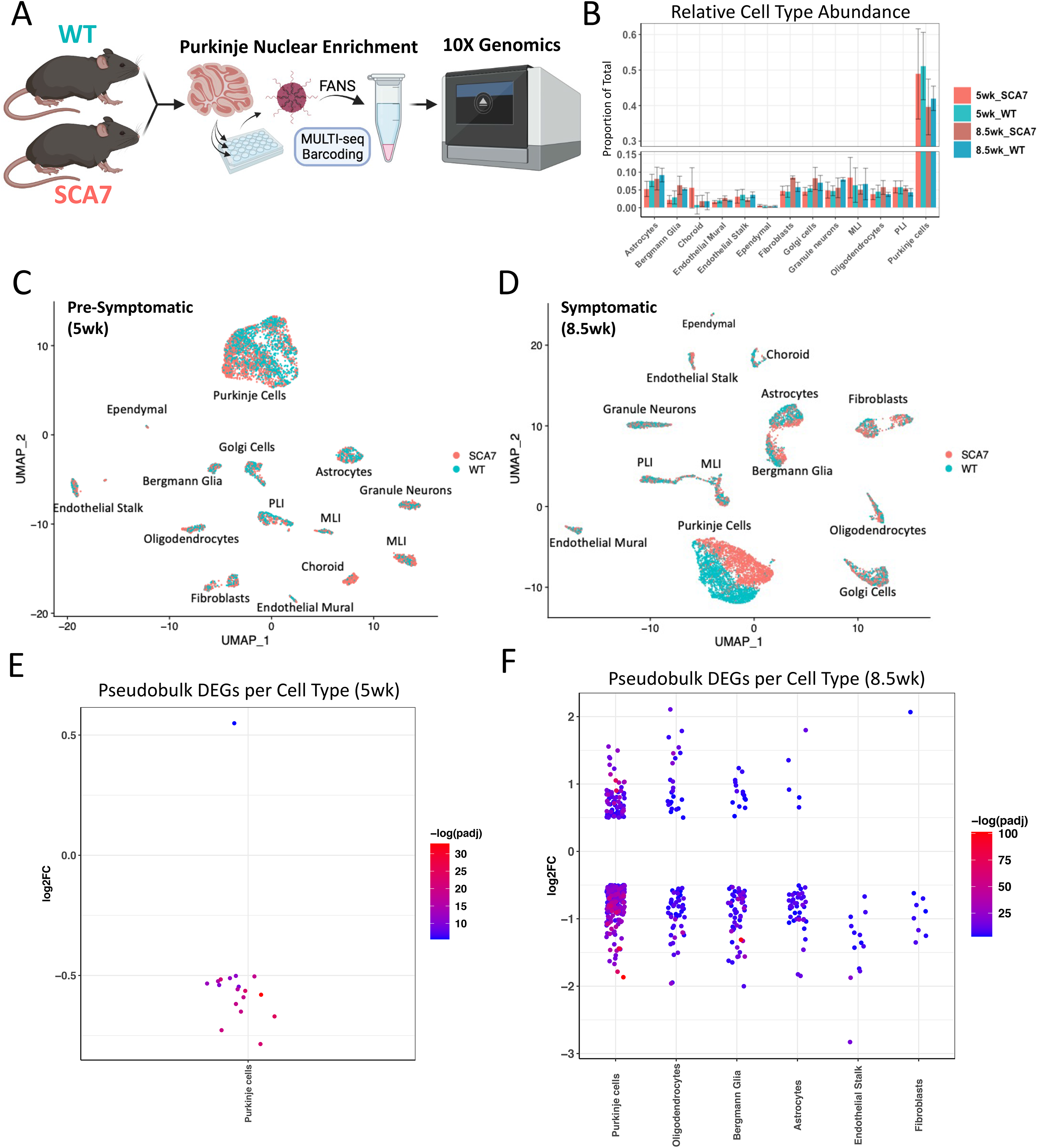
Purkinje-Enriched snRNA-seq Yields a High-Resolution Map of SCA7 Cerebellum. A) Experimental schematic detailing Purkinje-enriched snRNA-seq method combined with MULTI-seq CMO barcoding for multiplexing biological replicates into a single 10X GEM well. B) Relative cell type abundance ratios quantified from snRNA-seq samples. C) UMAP dimensional reduction of 5 week-old pre-symptomatic Purkinje-enriched snRNA-seq data with points colored by genotype of origin (coral = SCA7; teal = WT). D) UMAP dimensional reduction for 8.5 week-old symptomatic timepoint. E) Differential Expression results per cell type between SCA7 and WT nuclei for the 5 week timepoint using replicate-aware pseudobulk DEseq2. F) Differential Expression results per cell type for 8.5 week timepoint. Each point represents a significant DEG (cutoffs: adjusted p_val < 0.05; |log2-fold change| > 0.5)

### Analysis of snRNA-seq DEGs confirms glial contribution to SCA7 due to loss of glial identity

Given the known contribution of *Gfa2*-lineage cell types to the non-autonomous Purkinje cell degeneration in SCA7^29^, we first wanted to investigate differential expression results for signs of Glial dysfunction. Indeed, across the three most frequent glial cell types we noticed a clear separation of WT and SCA7 UMAP sub-clusters (**Figure 2A**) with a striking downregulation of key glial identity genes represented among SCA7 DEGs: oligodendrocytes (*Mobp*, *Qk*, *Plp1*), Bergmann glia (*Slc1a3* [EAAT1], *Qk, Sparcl1*), and astrocytes (*Aqp4*, *Sparcl1*, *Slc1a2* [EAAT2]) (**Figure 2B**). Further analysis suggested that the coordinate downregulation of glial identity genes in SCA7 cell clusters may stem from significantly decreased RNA expression of the *Qk* gene, which encodes the Quaking RNA binding protein (QKI). Loss of function of *Qk* is the genetic basis of the *quaking* mutant mouse, which shows severe myelination defects leading to ataxia, tremor, and early death^39,40^. QKI is a critical regulator of glial maturation and function through binding and stabilization of glial-specific transcripts^41^. Indeed, SCA7 DEGs from glial clusters show a highly significant overlap with QKI target mRNAs recently discovered by CLIP-seq^42^ (**Figure 2C**). Our snRNA-seq results thus corroborate previous reports of a glial contribution to SCA7^29^, and highlights one mechanistic hypothesis for the glial dysfunction in SCA7 through the impaired activity of QKI.

**Figure 2:**
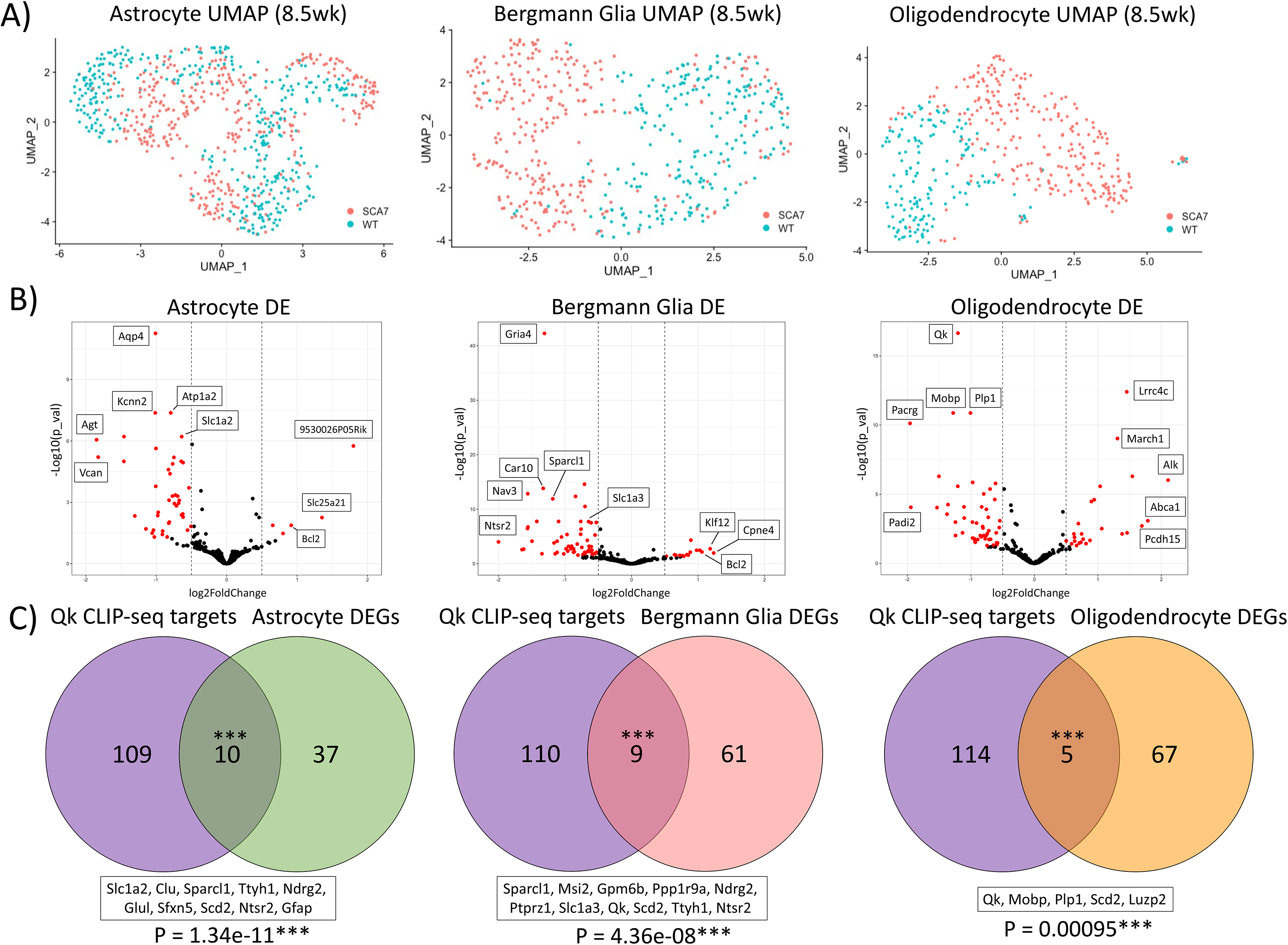
Signatures of Glial Identity Loss Identified in SCA7-266Q Animals. A) UMAP dimensional reduction for Astrocytes, Bergmann Glia, and Oligodendrocytes at the 8.5 week time point after re-clustering. Points colored for genotype of origin (coral = SCA7; teal = WT). B) Volcano Plots showing differential expression results between SCA7 and WT nuclei in Astrocytes, Bergmann Glia, and Oligodendrocytes. Log2FC and –log_10_(p-value) for each gene is shown with significant DEGs in red (cutoffs: adjusted p_val < 0.05; |log2-fold change| > 0.5). D) Venn-Diagram showing overlap of significant DEGs from each glial cell type with CLIP-seq targets of the QK RNA-binding protein. Hypergeometric test used to calculate overlap enrichment. Bonferroni adjusted p-value for overlaps shown along with Gene IDs.

### Purkinje cell DEGs highlight altered inhibitory synapse organization in SCA7 cerebellum

When we considered the full snRNA-seq dataset, we observed that PCs contained the greatest transcriptional changes, with 18 DEGs in presymptomatic mice and 320 DEGs in symptomatic animals. Sub-setting and re-clustering of PC nuclei revealed a dramatic progression of SCA7 transcriptional dysregulation from 5 weeks to 8.5 weeks of age (**Figure 3A-D**), corresponding to the timing of motor symptom onset. To gain further insight into the biological impact of the progressively dysregulated SCA7 transcriptome, we performed Gene Ontology (GO) analysis to categorize the Purkinje cell DEGs by their biological process. GO analysis revealed an enrichment of genes involved with Synaptic Signaling and Synapse Organization (**Figure 3E**). Many cell-adhesion molecules were represented in this biological category (*Lrrc4c*, *Cntn5, Grid2ip*, *Homer3*), suggesting a remodeling of SCA7 PC synaptic inputs upon disease onset (**Supplementary Table 2**). However, upon deeper analysis we determined that this enrichment of synaptic organization genes may be driven by a broader interference to PC subtype regulation.

**Figure 3:**
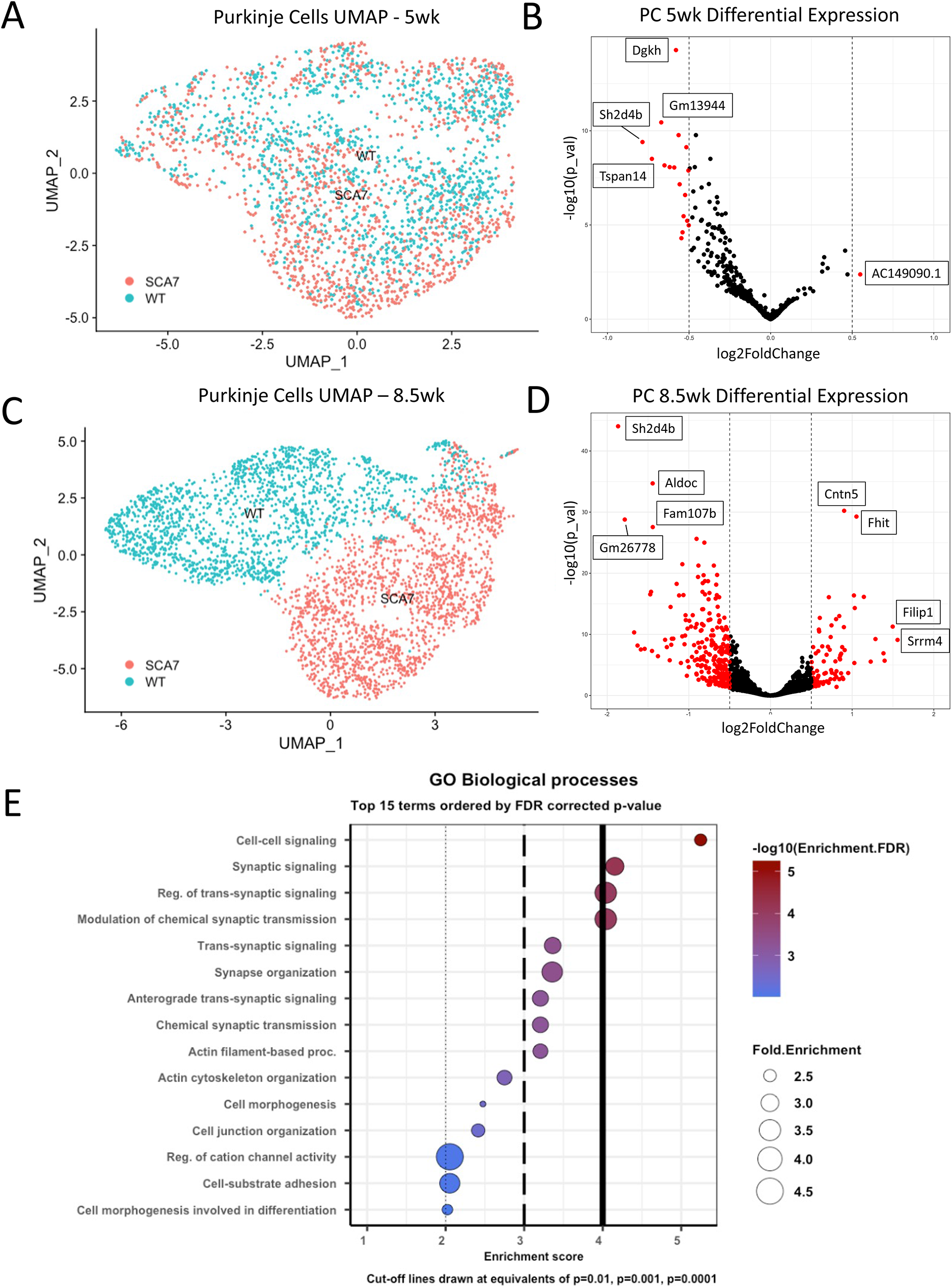
Progressive Transcriptional Dysregulation in SCA7 Purkinje Cells Highlights Changes in Synaptic Signaling. A) UMAP dimensional reduction for Purkinje Cell nuclei after subsetting and re-clustering at 5 week time point. B) Volcano Plot showing differential expression results between presymptomatic SCA7 and WT Purkinje nuclei with pseudobulk DEseq2. Significant DEGs shown in red (cutoffs: adjusted p_val < 0.05; |log2-fold change| > 0.5). C) UMAP dimensional reduction for Purkinje Cell nuclei after subsetting and re-clustering at 8.5 week time point. D) Volcano Plot showing differential expression results between symptomatic SCA7 and WT Purkinje nuclei with pseudobulk DEseq2. Significant DEGs in red (cutoffs: adjusted p_val < 0.05; |log2-fold change| > 0.5). E) Gene Ontology categories for 8.5 week symptomatic DEGs (320 DEGs) calculated with ShinyGO v0.77. Biological Process categories are sorted by -Log10(FDR) with vertical cutoff lines drawn at equivalent adjusted p-values of (dotted: p=0.01, dashed: p=0.001, solid: p=0.0001).

### The transcriptional differentiation of zebrin-II subtypes is severely diminished in the cerebellum of SCA7 mice

It has long been appreciated that the cerebellum of higher organisms, ranging from bony fish to mammals, exhibits an alternating pattern of gene expression of aldolase-C (*Aldoc)*, which upon immunostaining yields a distinct striping pattern, accounting for why aldolase-C came to also be known as ‘zebrin-II’^32,43^. Numerous studies of the cerebellum have established that this gene expression patterning of zebrin-II positive PCs and zebrin-II negative PCs organizes the cerebellum into distinct functional zones, with divergent cellular morphology, electrophysiological properties, and circuitry^33,44^. When one group recently compared the transcriptomes of zebrin-II positive and negative PCs in mouse cerebellum, they demonstrated that zebrin-negative PCs are specialized for synaptic plasticity during motor learning^45^.

To determine if the transcriptional dysregulation occurring in SCA7 PC degeneration affects zebrin-II transcriptional signatures, we investigated sub-clusters of PC nuclei from SCA7 knock-in mice and WT mice in our snRNA-seq gene expression data. UMAP dimensional reduction at the early symptomatic time point displayed a highly divergent axis of SCA7 nuclear profiles that strongly correlated with zebrin-II expression, and revealed a marked decrease in the fraction of zebrin-II positive PCs (**Figure 4A**). To better assess the status of zebrin-II transcriptional signatures in SCA7 PCs, we initially sought to define a set of zebrin-II identity genes from WT PCs alone. Sub-clustering of WT PCs revealed a clear demarcation of the zebrin-II transcriptional signature, with UMAP clusters separating zebrin-II positive PCs from zebrin-II negative PCs (**Figure 4B**). When we performed differential expression analysis of the WT zebrin-II PC subtypes, we identified 354 significant DEGs that accounted for the demarcation of zebrin-II positive and zebrin-II negative PCs in WT mouse cerebellum (**Figure 4C**). Strikingly, these WT zebrin-II axis genes showed highly significant overlap with SCA7 PC DEGs (*P* < 1 x 10^-30^), with 21% of SCA7 PC DEGs now identifiable as zebrin-II identity genes (**Figure 4D**). Treating this ‘Zebrin-II axis’ list as a biological category and then comparing FDR enrichment to the GO terms assessed in **Figure 3E** confirmed that the impact of SCA7 transcription dysregulation to the ‘Zebrin-II-axis’ genes far exceeds the highly significant effect on the Synaptic signaling genes (**Figure 4E**).

**Figure 4:**
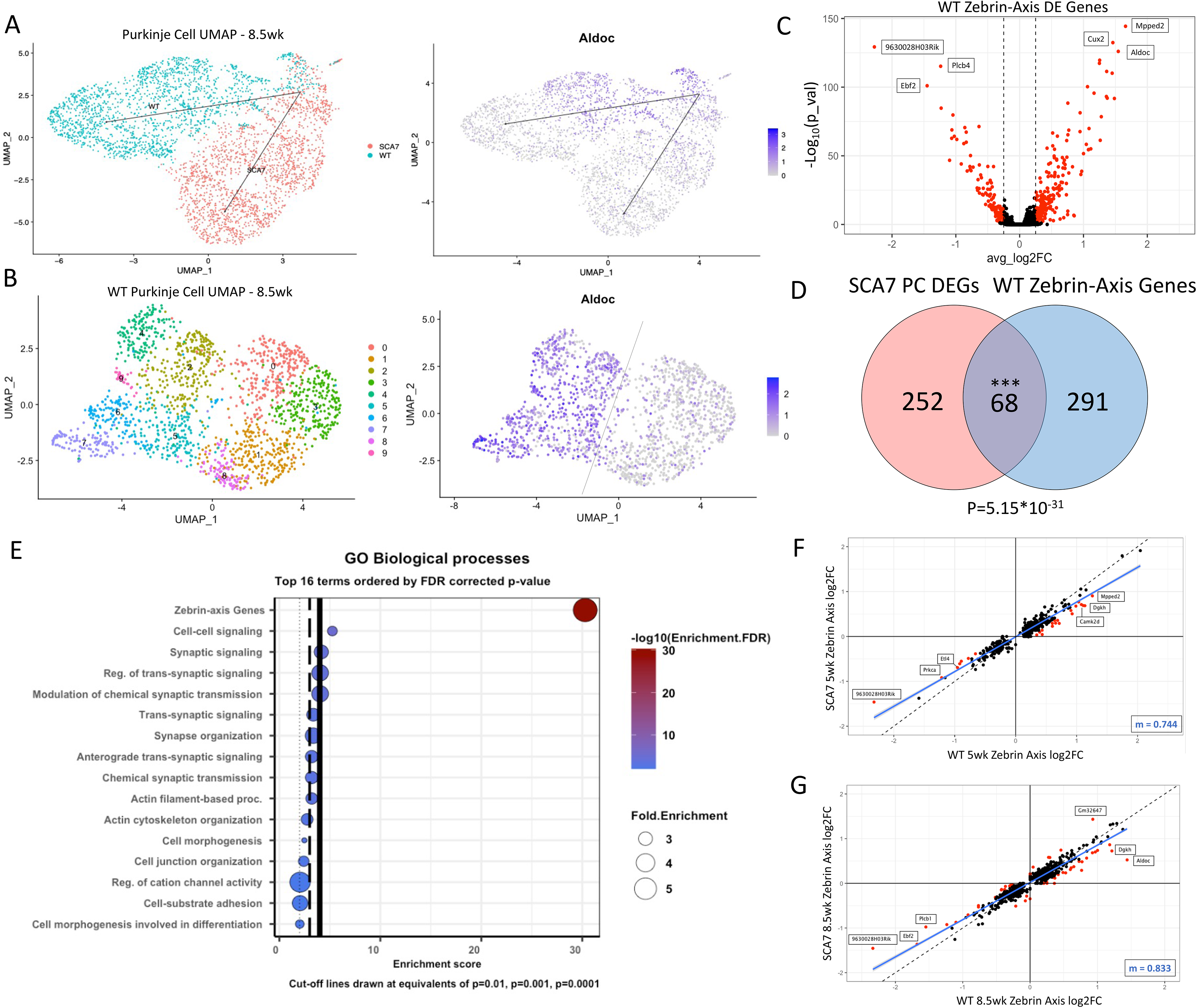
Zebrin-II Subtype Dysregulation Explains Highest Proportion of SCA7 Purkinje Cell DEGs. A) UMAP dimensional reduction for 8.5 week-old Purkinje Cells shown alongside normalized Aldoc (Zebrin-II) expression for each single-nucleus. B) WT Purkinje Cells subset and re-clustered to find Zebrin-positive and Zebrin-negative clusters. C) Volcano Plot showing differential expression between WT Zebrin-positive and Zebrin-negative clusters. Wilcoxon Rank Sum test was used and significant genes are shown in red (cutoffs: adjusted p_val < 0.05; |log2FC| > 0.25). D) Venn-Diagram showing overlap of 8.5 week-old SCA7 DEGs in PCs with Significant Genes of the WT Zebrin-Axis. Statistical enrichment calculated with Hypergeometric test and adjusted with Bonferroni correction of 13,078 representing number of Gene Ontology Biological Process Categories (adjusted p_val = 5.15e-31***). E) Gene Ontology Biological Processes from Figure 3B re-plotted with Zebrin-Axis gene category added. F) WT Zebrin-axis DEGs calculated from 5 week-old PCs (as in panel B) and plotted against 5 week-old SCA7 Zebrin-axis DEGs by Log2FC value. Linear Regression was applied to find correlation of Log2FC values from WT and SCA7 (slope = 0.744). Points are colored red if Log2FC difference between WT and SCA7 is greater than 0.25. G) WT and SCA7 Zebrin-axis comparative plot for 8.5 week-old PCs (slope = 0.833).

We then assessed the transcriptional dysregulation of zebrin-II PC subtypes in SCA7 by sub-clustering SCA7 PCs alone and computing differential expression between clusters of zebrin-II positive and zebrin-II negative PCs. The SCA7 PC zebrin-II-axis showed significant abnormalities when compared to WT PC zebrin-II-axis in both presymptomatic and symptomatic mice. Indeed, in 5 week-old SCA7 mice, the zebrin-II-axis gene expression signatures already display a clear trend toward the direction of zebrin identity loss, with log-fold expression of zebrin-II identity genes dampened in both the zebrin-II positive and zebrin-II negative PC subtypes (**Figure 4F**). When we plotted the log-fold expression values of zebrin-II axis genes for WT vs. SCA7 PCs and performed linear regression to quantify this trend, we calculated a decreased slope (m = 0.774) which approximates a line plotted with a slope for a theoretically perfect correlation (**Figure 4F**). This gene expression signature in presymptomatic SCA7 mice indicates that the presence of polyQ-expanded ataxin-7 in PC nuclei is likely interfering with the regulatory mechanisms that specify zebrin-II PC subtypes prior to the appearance of visible nuclear aggregates. Symptomatic 8.5 week-old SCA7 mice display progressive dysregulation of the zebrin-II axis, with the emergence of numerous significantly up- and down-regulated zebrin-II-identity genes, with even greater disruption of zebrin-II identity (**Figure 4G**).

### Disruption of zebrin-II striping in symptomatic SCA7 mice and in related polyQ SCAs

Given the marked downregulation of the canonical zebrin-II marker gene *Aldoc* in the PCs of symptomatic SCA7 mice (**Figure 4C**), we hypothesized that the stereotypical parasagittal striping pattern of the cerebellum might be affected. To investigate the spatial expression pattern of zebrin-II, we performed immunohistochemistry on coronal cerebellar sections obtained from SCA7 266Q knock-in mice and WT littermates. Although immunostaining of zebrin-II in 5 week-old presymptomatic SCA7 mice revealed normal patterning with robust striped expression in the posterior lobules, we documented a dramatic, near-complete loss of the zebrin-II striping pattern in 8.5 week-old SCA7 mice, despite limited evidence of PC degeneration based upon robust calbindin-1 expression (**Figure 5A**). To quantify this phenotype, we developed an image analysis pipeline to compute the amplitude of zebrin-II stripe expression across lobule VIII of the cerebellum, and verified that zebrin-II striping integrity is virtually absent by this time point in SCA7 mice (**Figure 5B**).

**Figure 5:**
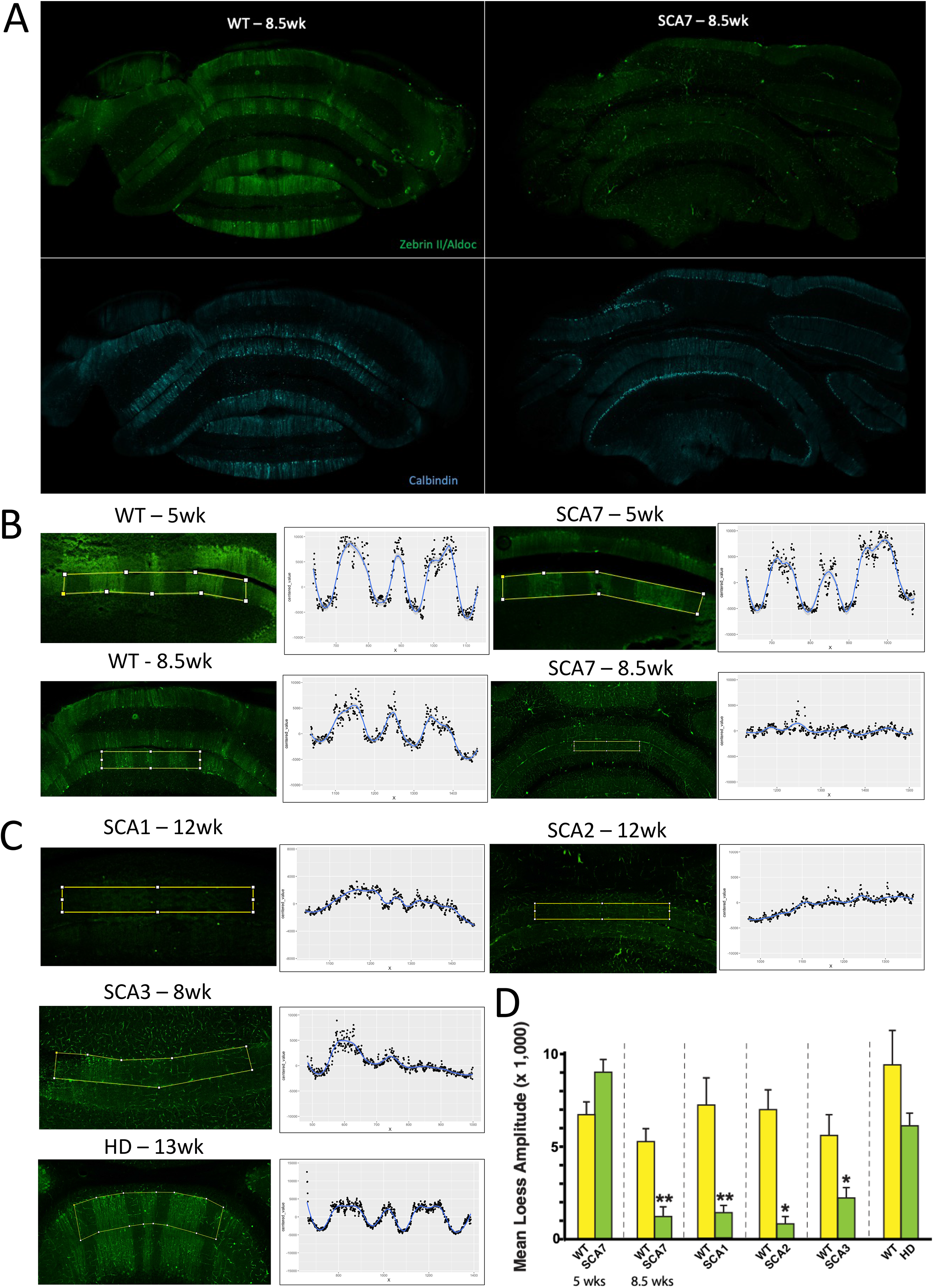
Zebrin-II Parasagittal Stripes are Lost in SCA7 and Related Polyglutamine Ataxias. A) Immunohistochemistry performed on coronal cerebellar slices from 8.5 week-old WT and SCA7 mice stained for Zebrin-II/Aldoc (green) and Calbindin (blue). B-C) Representative images of Zebrin staining intensity across the central 3 stripes of Cerebellar Lobule VIII paired with median pixel intensity values across the measured area. Intensity values were fit with a Loess Polynomial Regression, before measuring the signal amplitude of each stripe and averaging to find the mean Zebrin intensity for each animal. Models and ages tested: WT 5 weeks (n=3), SCA7 5 weeks (n=3), WT 8.5 weeks (n=3), SCA7 8.5 weeks (n=3), SCA1-175Q 12 weeks (n=3), SCA2-127Q 12 weeks (n=2), SCA3-YAC (Q84/Q84) 8 weeks (n=4), and HD-N171-82Q 13 weeks (n=3). Note that age-matched WT littermates for SCA1, SCA2 and HD models were also analyzed for Zebrin stripe intensity. D) Amplitudes for each group were compared to respective WT controls with a one-tailed T-test. (SCA7 8.5 weeks, **p = 0.0065; SCA1-175Q 12 weeks, **p= 0.0083; SCA2-127Q 12 weeks, *p= 0.016, SCA3-YAC-84Q, *p=0.049). Error bars = s.e.m.

A number of studies have noted striking similarities between the transcriptional alterations in the cerebellum of various polyQ SCAs, correlating findings from SCA1, SCA2, and SCA7^24,46^. To determine if this shared transcriptional pathology also impacts the normal developmentally programmed parasagittal striping of the cerebellum, we examined zebrin-II spatial expression in the cerebellum of SCA1 175Q knock-in mice and SCA2 127Q transgenic mice, two extensively studied mouse models believed to be representative of their cognate human diseases^4,5^. Zebrin-II immunostaining of coronal cerebellar sections similarly revealed a complete ablation of the normal parasagittal striping pattern in both early symptomatic SCA1 and SCA2 mice (**Figure 5C**). We also observed a significant disruption of parasagittal striping in early symptomatic SCA3-YAC-Q84 mice, another well studied polyQ SCA model^6,47^ (**Figure 5C-D**). To determine if this phenomenon also occurs in Huntington’s disease (HD), which is characterized by high-level cerebellar expression of polyQ protein – but without cerebellar degeneration, we then performed zebrin-II immunostaining of coronal cerebellar sections from a well-known HD transgenic mouse model – N171-82Q^48^, and remarkably we observed a perfectly normal parasagittal striping pattern in the cerebellum of early symptomatic HD N171-82Q mice (**Figure 5C-D**). These results indicate that disrupted zebrin-II identity is a shared feature of polyQ SCAs.

### Human SCA7 Cerebellum displays Zebrin-II Dysregulation in Molecular Layer Interneurons

While the presence of disrupted zebrin-II expression patterning in SCA7 model mice is a compelling finding, an important question is whether this phenomenon is relevant to the disease process in human patients. To address this issue, we performed multiplexed snRNA-seq on post-mortem cerebellum samples obtained from five SCA7 patients and four unaffected controls, and applied our Purkinje-enriched nuclear isolation method and MULTI-seq barcoding as described earlier. When we analyzed the human snRNA-seq data using a standard Seurat normalization and clustering pipeline, we were unable to detect any PC nuclei based upon cell type identification of UMAP clusters by marker gene expression (**Figure 6A**). As it turns out, this was not unexpected, as several studies of post-mortem human cerebellum have encountered this identical technical issue, and have failed to recover PC nuclei to date^49,50^. However, when we reviewed the UMAP clustering of our human cerebellum snRNA-seq results, we noted significant enrichment of both Type-I and Type-II Molecular Layer Interneurons (MLI), which revealed significant gene expression alterations between SCA7 patients and unaffected controls (**Figure 6B**). Similar to our observation in SCA7-266Q mice, SCA7 human tissues showed a dramatic cell type selective decrease in RNA abundance, with outsized impacts on MLIs (**Supplemental Figure S4**). Looking at gene expression changes in SCA7 MLIs, *ALDOC* (zebrin-II) displayed a significant downregulation in both MLI-1 and MLI-2 clusters, confirming that marked dysregulation of zebrin-II gene expression is a shared feature of the disease process in the cerebellum of human SCA7 patients (**Figure 6B-C**).

**Figure 6:**
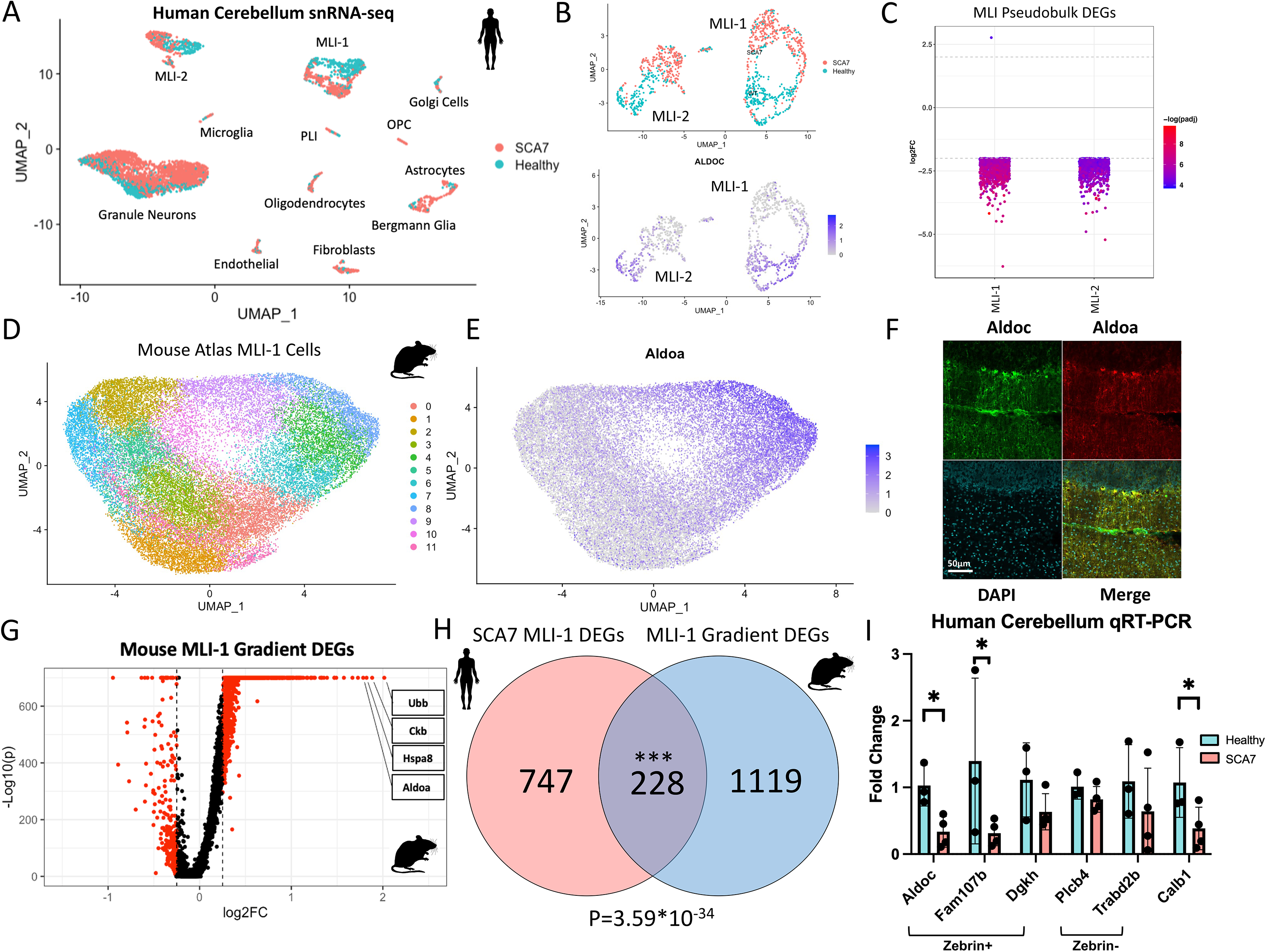
Human SCA7 Cerebellar Tissues Show Degradation of Zebrin-Related Subtypes in Molecular Layer Interneurons. A) UMAP clusters for snRNA-seq performed on Human post-mortem cerebellar tissues from 5 SCA7 patients and 4 unaffected controls colored by genotype (coral = SCA7; teal = WT). B) Molecular Layer Interneuron sub-clustering results showing strong co-clustering of SCA7 nuclei, and strong down-regulation of Zebrin-II/ALDOC. Panels are colored by Disease Status (top) and normalized ALDOC expression (bottom). C) Differential expression results comparing SCA7 patients to unaffected controls for each recovered cell type using pseudobulk DEseq2. (Cutoffs: adjusted p_val < 0.05; |log2-fold change| > 2). D) UMAP sub-clustering of MLI-1 cells from the Mouse Cerebellar Atlas colored by UMAP cluster. E) Strong gradient of normalized *Aldoa* expression in Mouse Atlas MLI-1 cells. F) IHC representative image from 5 week-old WT mouse cerebellum showing striped expression of Aldoc along with Aldoa. G) Volcano plot differential expression results using Wilcoxon Rank Sum test between Aldoc/Aldoa-high and Aldoc/Aldoa-low clusters from the Mouse Cerebellar Atlas (cutoffs: adjusted p_val < 0.05; |log2-fold change| > 0.25). H) Venn-Diagram showing overlap of SCA7 Human MLI-1 DEGs with Mouse MLI-1 Aldoc/Aldoa Gradient DEGs. Statistical enrichment of overlap computed with hypergeometric test shown with Bonferroni corrected p-value (p=3.59e-34). I) qRT-PCR for Zebrin Identity genes on bulk RNA extracted from human SCA7 patients (n=4) and unaffected control (n=3) cerebella. Fold change shown relative to mean Ct value for control samples; statistics calculated with one-tailed t-test.

Our detection of altered zebrin-II expression in human SCA7 MLIs raises a provocative question: Are MLIs also subject to a developmental program of alternating gene expression linked to PC parasagittal striping? Indeed, a number of previous studies have suggested that both the morphological and electrophysiological properties of MLIs appear bounded to parasagittal zones, generating functional modular circuits with zebrin-II positive and negative PCs^51–53^. To definitively evaluate the gene expression properties of MLIs, we reanalyzed MLIs from the Kozareva et.al. Transcriptomic Mouse Cerebellar Atlas^25^ to determine if zebrin-II and its co-regulated genes display a gene expression gradient akin to PCs (**Figure 6D**). This analysis revealed that both MLI-1 nuclei and MLI-2 nuclei from the Mouse Cerebellar Atlas form robust subclusters with a detectable zebrin-II gene expression gradient, marked even more prominently by *Aldolase-a* (*Aldoa*) expression (**Figure 6E, Supplemental Figure S5**). To assess the biological relevance of this expression gradient, we immunostained mouse cerebellar sections for *Aldolase-*a and confirmed the existence of an *Aldolase-a* striping pattern that clearly parallels the alternating expression pattern of zebrin-II (**Figure 6G**). Using subclusters with ‘high’ and ‘low’ expression of *Aldoc/Aldoa*, we performed differential expression analysis to delineate the entire list of genes defining this gradient. This analysis yielded 1,347 significant DEGs for MLI-1 and 1,437 significant DEGs for MLI-2 nuclei, with highly overlapping gene sets that form a gene expression gradient correlated with *Aldoc* (**Figure 6G, Supplemental Figure S5**). To evaluate the relationship between the MLI gene expression changes detected in human SCA7 patients and MLI gene expression patterning detected in mouse cerebellum, we compared the human SCA7 MLI-1 DEGs with the DEGs defining mouse MLI-1 Zebrin subclusters, and we noted an extremely high degree of enrichment (*P* = 3.59 x 10^-34^), indicating that MLI-1 patterning genes are disproportionally impacted in SCA7 patients (**Figure 6F**). Finally, to examine the expression level of PC zebrin-II identity genes in human SCA7 patient postmortem cerebellum, we pursued qRT-PCR analysis on bulk cerebellar RNA (in lieu of PCs in single-cell RNA sequencing), and we confirmed significant downregulation of PC zebrin-II identity genes in SCA7 patients (**Figure 6H**).

### PC inhibitory synapse activity is increased in the cerebellum of SCA7 model mice

Although changes in the expression of zebrin-II subtype genes and the resulting disappearance of zebrin-II parasagittal stripes in SCA7 model mice are remarkable, an important question is how these findings alter cerebellar function and thereby contribute to SCA7 cerebellar degeneration. Since the Gene Ontology Biological Function categories most dysregulated in SCA7 PCs are overwhelmingly related to Synapse Organization (**Figure 3E**), and Zebrin-II subtypes are known to differ in synaptic functions and electrophysiology^33^, we hypothesized that the breakdown of Zebrin subtype maintenance would have downstream consequences on PC synapses. To address this hypothesis, we assessed PC synapse organization by performing quantitative immunohistochemistry of both excitatory and inhibitory synapses in the cerebellar molecular layer. Purkinje cell dendrites receive synaptic input through three distinct synapses marked by the presynaptic vesicular proteins VGAT (inhibitory)^54^, VGLUT1 (excitatory)^55^, and VGLUT2 (excitatory)^56^. VGLUT1 marks excitatory synapses originating from the highly dense parallel fibers, which emanate from granule cell neurons and elicit high frequency simple spikes in PCs^57,58^, while VGLUT2 marks excitatory synapses formed by the climbing fibers, which originate from the inferior olive and show a tight 1:1 relationship with PCs^59^. Climbing fiber synapses operate at a lower frequency and generate the complex spikes in PCs^60^. We stained WT and SCA7 cerebella with antibodies against VGLUT2 paired with calbindin-1 to mark PC dendrites and quantified synaptic puncta in lobule VII of the cerebellar vermis. At both the presymptomatic and symptomatic stages, climbing fiber synapses in SCA7 mice appeared normal, with VGLUT2 synaptic density and innervation along the length of the PC dendrite matching what is seen in WT mice (**Supplemental Figure S6A-D**), in agreement with a previous report^24^. To profile inhibitory synapses, we stained for presynaptic VGAT and postsynaptic Gephyrin proteins, and performed calbindin-1 co-staining to differentiate inhibitory synapses on PC dendrites from synapses on the surrounding molecular layer inhibitory interneurons (MLI). In 5 week-old pre-symptomatic SCA7 mice, immunostaining quantification indicated a similar number of both PC and non-PC inhibitory synapses when compared to 5 week-old WT littermate controls, though there was a trend toward increased numbers of inhibitory synapses in SCA7 cerebellum (**Figure 7A-B**). In 8.5 week-old symptomatic SCA7 mice, however, we observed a marked increase in the number of inhibitory synapses – both on PC dendrites and on surrounding MLI (**Figure 7C-D**). These findings confirm that PC gene expression changes observed by snRNA-Seq result in neuroanatomical alterations in cerebellum synapse organization in SCA7 model mice.

**Figure 7:**
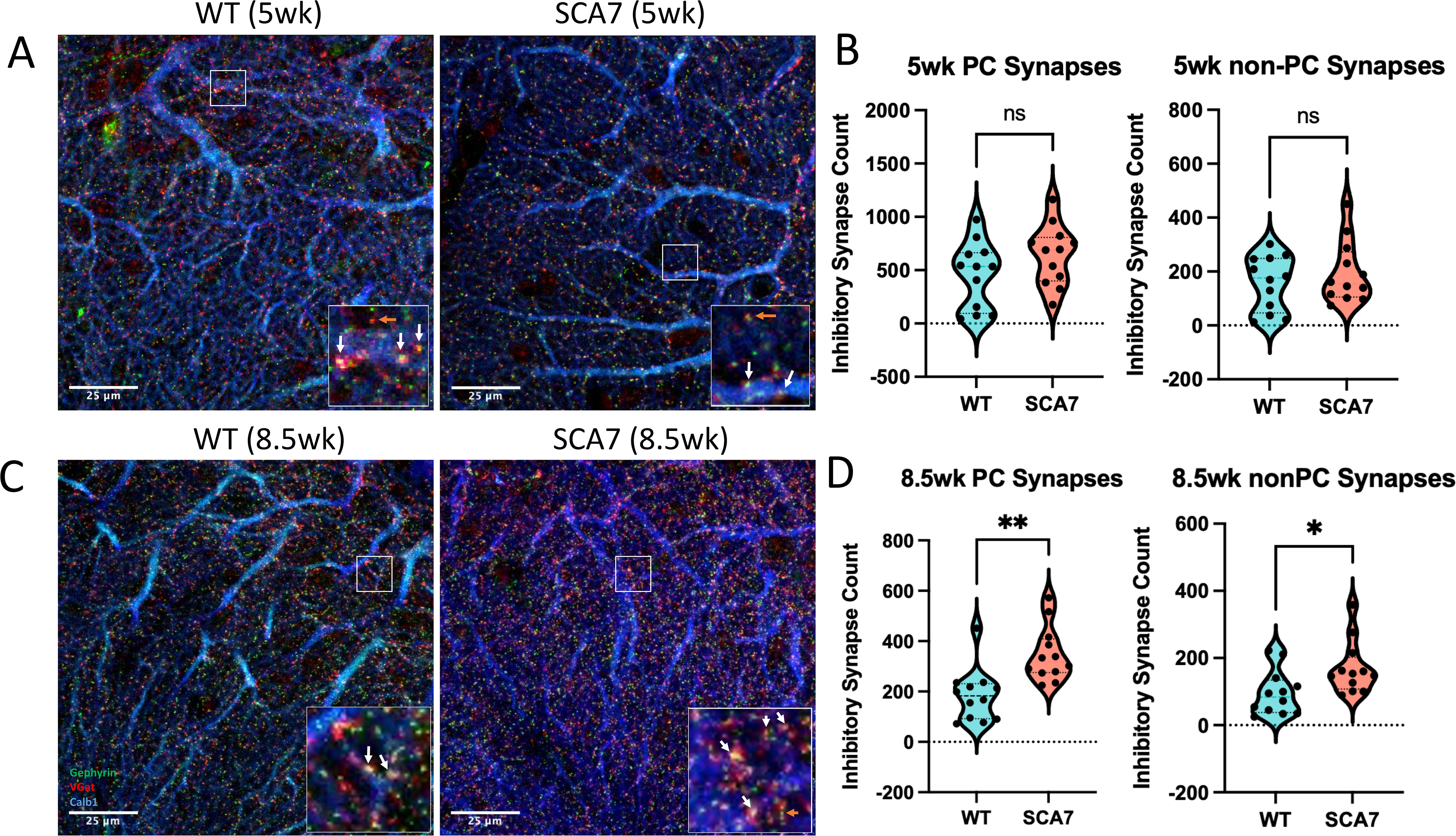
Symptomatic SCA7-266Q Mice Contain Increased Inhibitory Synapses. A) Inhibitory Synapse immunohistochemistry performed on the molecular layer of Lobule VII in 5 week-old WT and SCA7-266Q mice (63X magnification; green = VGAT, red = Gephyrin, blue = Calbindin-1). Image in-sets contain arrows pointing to yellow co-localized signal of intact inhibitory synapses on Calbindin-1 positive PC dendrites. Orange arrow demarcates a non-PC synapse. B) Quantification of inhibitory synapses by intact synapse numbers in a 15 μm z-scan. Inhibitory synapses overlapping Calbindin-1 were classified as PC-synapses. n=6 mice/genotype; two images per animal; statistics calculated with two-tailed t-test: PC Synapses, p=n.s., non-PC Synapses, p=n.s. C) Inhibitory Synapse immunohistochemistry performed on the molecular layer of Lobule VII in 8.5 week-old WT and SCA7-266Q mice (63X magnification; green = VGAT, red = Gephyrin, blue = Calbindin-1). Image in-sets contain arrows pointing to yellow co-localized signal of intact inhibitory synapses on Calbindin-1 positive PC dendrites. Orange arrow demarcates a non-PC synapse. D) Quantification of inhibitory synapses by intact synapse numbers in a 15 μm z-scan. Inhibitory synapses overlapping Calbindin-1 were classified as PC-synapses. n=6 mice/genotype; two images per animal; statistics calculated with two-tailed t-test: PC Synapses **p=0.001, non-PC Synapses *p=0.021.

To determine the functional consequences of altered inhibitory synapse organization in symptomatic SCA7 mice, we directly examined Purkinje neuron electrical activity in acute cerebellar slices. Unlike previous studies that assessed the intrinsic pace-making activity of PCs and documented evidence of irregular spiking^24,61^, here we focused on local circuits upstream of PC firing by performing whole-cell voltage clamp recordings to measure spontaneous inhibitory post-synaptic currents (sIPSCs). Analysis of sIPSCs uncovered significantly increased inhibitory synaptic amplitudes in SCA7 PCs from 8.5 week-old mice (**Figure 8A**), with no change in sIPSC frequency compared to WT littermates. Excitatory post-synaptic currents (sEPSCs) in SCA7 PCs displayed normal amplitudes and decreased sEPSC frequency (**Figure 8B**). We then collected cell-attached recordings of PC action potentials and quantified the spiking rate and regularity through inter-event-intervals. SCA7 PCs displayed a significant decrease in spiking rates (**Figure 8C**), reflecting the shift in excitatory/inhibitory synaptic balance on the PC dendrites.

**Figure 8:**
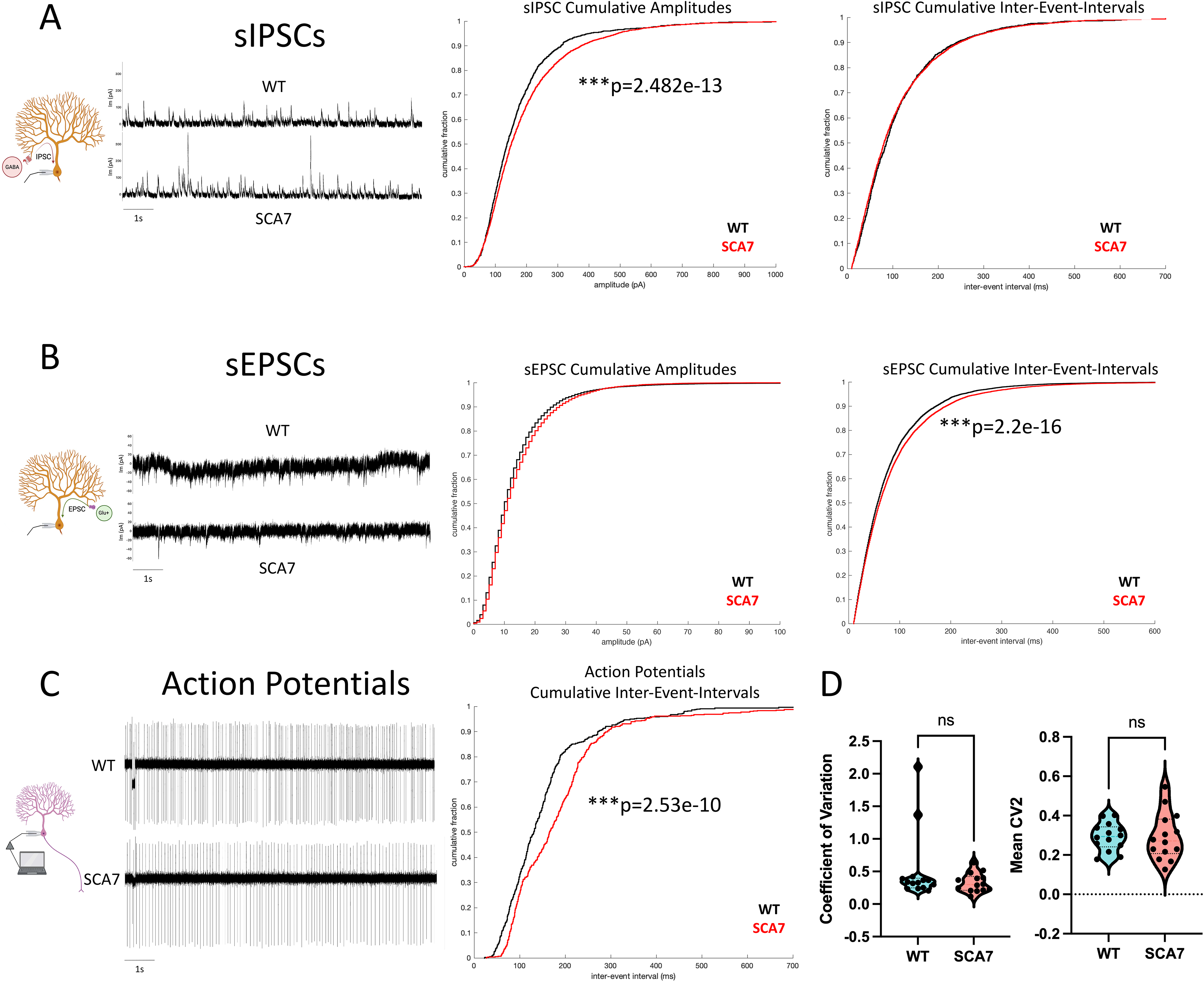
Increased Inhibitory Post-Synaptic Currents in SCA7 Purkinje Cells Leads to Reduced Spiking Rates. A) Spontaneous Inhibitory Post-Synaptic Currents (sIPSC) recorded from Purkinje cells in acute cerebellar slices from 8.5 week-old SCA7-266Q mice and matched WT littermates. Whole-cell voltage-clamped sIPSCs were measured at +10mV for n=6 WT and n=15 SCA7 Purkinje Cells. Amplitudes and Inter-Event-Intervals were measured for each sIPSC event before randomly sampling an equal number from each cell, shown plotted in a cumulative histogram. Wilcoxson Rank Sum test for IPSC amplitudes, ***p=2.48×10^-13^. B) Spontaneous Excitatory Post-Synaptic Currents (sEPSC) recorded from Purkinje cells in acute cerebellar slices from 8.5 week-old SCA7-266Q mice and matched WT littermates. Whole cell voltage-clamped sEPSCs were measured at -75mV for n=18 WT and n=17 SCA7 Purkinje Cells. Amplitudes and Inter-Event-Intervals were measured for each sEPSC and randomly sampled, shown plotted in a cumulative histogram. Wilcoxson Rank Sum test for EPSC inter-event-intervals, ***p=2.20×10^-16^. C) Cell-attached recordings were taken from n=14 WT and n=14 SCA7 Purkinje Cells. Action Potentials were analyzed to find inter-event-intervals and randomly sampled from each cell to produce a cumulative histogram. Wilcoxon Rank Sum test for Action Potential Inter-Event-Intervals, ***p=2.53×10^-10^. D) The Coefficient of Variation in recorded action potentials was measured for each cell, as well as the CV2 statistic. No significant differences in spiking regularity were found.

## DISCUSSION

We, and others, have identified transcription dysregulation as a central feature of SCA7 disease pathogenesis, stemming from the realization that ataxin-7 is a core component of the STAGA co-activator complex, and that mutant polyQ-ataxin-7 protein gets incorporated into the STAGA complex and therein disrupts the function of transcription factors that depend upon STAGA for transactivation of their target genes^13,20^. We previously performed bulk RNA-seq analysis on cerebellum from presymptomatic and early symptomatic SCA7 mice, and documented a role for altered calcium regulation in SCA7 cerebellar degeneration, linking the calcium dysregulation to dysfunction of Sirtuin-1, a master regulator of gene expression and cellular function^61^. While this work yielded key insights into the molecular basis of SCA7, there are two major limitations in the biological interpretation of such bulk RNA-seq datasets: 1) RNA reads cannot be assigned to a particular cell type, and 2) the cerebellum is dominated by the abundant granule cell neurons (GCNs), as GCNs comprise >90% of cells in the cerebellum and thereby dilute signals from non-GCN cell types. As Purkinje cell (PC) neurons integrate neural circuit information to provide critical output to the deep cerebellar nuclei and thence to the entire motor system of the brain, and PCs undergo degeneration in SCA7, evaluation of PC gene expression is crucial to obtain a thorough understanding of the SCA7 disease process. Here we introduced use of a PC nuclear enrichment protocol, and successfully applied it to perform snRNA-seq to interrogate cerebellar degeneration in SCA7 266Q knock-in mice. By evaluating both presymptomatic and early symptomatic SCA7 mice in comparison to age- and sex-matched WT littermate controls, we discovered alterations in PC gene expression leading to changes in synapse organization and zebrin-II patterning that likely underlie SCA7 disease pathogenesis.

The cerebellum is a complex brain structure that plays a critical role in motor coordination and learning, functionally organized into parasagittal modules marked by zebrin-II. While zebrin-II expression is typically appreciated as a marker of PC subtypes, these functional modules extend to the surrounding MLIs, which we have now characterized by the differential expression of over 1000 genes. While the significance of zebrin-II parasagittal expression striping is still not fully understood, it has been implicated in several functions, including the control of motor learning, organization of cerebellar circuitry, and regulation of synaptic plasticity^34,62^. The development of the cerebellum is an incredibly refined and meticulous process that extends well into the postnatal period before completion in humans and mice. PCs can be initially recognized in the developing mouse brain at embryonic day 10.5 (E10.5), with a second layer of PCs emerging at E12.5^63–66^. Based on single-cell studies, at least 4 “embryonic” clusters of PCs can be identified in the developing cerebellum with a specified set of marker genes unique to this developmental window^67^. Although SCA7 mice retain parasagittal striping of the cerebellum at 5 weeks of age, evidence for gene expression alterations in zebrin identity genes are already present at this time point (**Figure 4**). These findings indicate that altered development of the cerebellum is a defining feature of SCA7 disease, and thus contribute to an emerging view of polyglutamine neurodegeneration where aberrant development is critical for setting the stage for later disease pathogenesis. For example, in SCA1, the timing of onset of mutant ataxin-1 gene expression is a powerful determinant of Purkinje cell degeneration, as delaying the expression of polyQ-ataxin-1 from postnatal day 2 (P2) to postnatal day 10 (P10) markedly protects against later cerebellar degeneration, even if much higher levels of polyQ-ataxin-1 protein expression occur starting at P10^68^. Furthermore, recent studies of Huntington’s disease (HD), another CAG-polyQ disorder, indicate that defects in the development of the cortex can be detected in human patients and model mice, underscoring a likely role for altered neurodevelopment in HD pathogenesis^69^. Furthermore, these neuronal circuits rely on direct feedback and support from surrounding glial cell types whose roles are indispensable for proper synaptic transmission and plasticity^70^.

A common theme in our investigation of the SCA7 transcriptional landscape was the recurrent dysregulation of cell type identity genes in both neuronal and glial cell types. Perhaps this observation should be expected in the context of SCA7 and a mutated STAGA complex, considering the central role of epigenetic marks and transcriptional regulation in the process of cellular differentiation, which yields the wide variety of cell types necessary for a functioning CNS. Previous studies have identified the importance of glial dysfunction in SCA7 pathogenesis through the reduced expression of the Bergmann glia-specific glutamate transporter EAAT1 (*Slc1a3,* GLAST)^29^, which we have validated here and expanded to include several other crucial glial transporters in Bergmann glia and astrocytes. We had previously shown that reduced EAAT1 activity led to a reduced ability of SCA7 glia to reuptake glutamate^29^, which may be directly connected to the increased inhibitory synapse transmission identified in this study.

A key observation, only made possible by snRNA-seq, was the identification of a progressive breakdown in PC zebrin-II subtypes, following on this pattern of cell identity loss. While standard pathway analyses may have limited our interpretation of gene expression changes to synaptic pathways, a targeted analysis of PC subclusters revealed that disruption of zebrin-II subtype genes is the predominant signal, leading to our discovery of zebrin-II stripe loss in symptomatic animals. To determine if disintegration of zebrin-II expression patterning is restricted to SCA7 or is a feature of other polyglutamine diseases, we immunostained cerebellar sections from three closely related polyQ SCA mouse models and a representative HD mouse model, and we documented disruption of parasagittal striping in the cerebellum of early symptomatic SCA1, SCA2, and SCA3 mice, but observed normal preservation of parasagittal striping in the cerebellum of HD mice. Hence, it appears that the effect of polyQ disease proteins on zebrin-II parasagittal striping correlates with the cerebellar disease process, as HD mice do not display cerebellar degeneration^71^. Previous studies have emphasized that SCA1, SCA2, and SCA7 display considerable overlap in transcriptional alterations in the degenerating cerebellum in highly representative mouse models^24,72^. It is therefore tempting to speculate that the disruption of zebrin-II identity could be at the core of the transcriptional pathology underlying these polyQ SCAs. Dysregulation of zebrin-II identity genes in multiple polyQ SCAs suggests a common mechanism of pathogenesis and could serve as a potential therapeutic target for the treatment of these disorders. Future studies are needed to map the extent of overlap between altered zebrin-II identity genes in the various SCAs and determine the mechanistic basis for these shared abnormalities. Overall, these findings provide novel insights into the molecular mechanisms underlying polyQ ataxias and could have important implications for the development of new therapies for these devastating diseases.

To determine if the zebrin-II dysregulation observed in SCA7 mice is relevant to the disease process in human patients, we obtained post-mortem cerebellum from five SCA7 patients and four unaffected human controls, and performed snRNA-seq on nuclei isolated from these samples. Although we could not obtain PCs, which is the typical experience based upon all other published reports to date^49,50^, we did detect MLIs and noted prominent zebrin-II dysregulation in SCA7 patients, which led us to evaluate MLIs and document alternating expression patterns in this cerebellar cell type. When we directly measured the expression levels of zebrin-II identity genes in bulk RNA samples obtained from SCA7 and control human cerebella, we observed decreased expression of a subset of these marker genes in SCA7 patients. Together, these results from SCA7 human cerebellar tissues further support the notion that dysregulated zebrin identity could be one of the foremost driving pathological transcriptional mechanisms leading to neurodegeneration and neuron dysfunction in SCA7, and therefore could be considered a new pathological hallmark of polyQ SCAs.

Another striking observation emerging from the SCA7 mouse snRNA-seq dataset was the upregulated expression of cell adhesion molecules involved in inhibitory synapse formation. To determine if increased expression of synapse organization genes in SCA7 PCs was paralleled by functional changes of biological relevance, we sought to quantify synapses and examine synapse activity in SCA7 cerebellum. We documented that early symptomatic SCA7 mice display markedly increased numbers of inhibitory synapses in the molecular layer of the cerebellum, accompanied by significantly increased inhibitory synaptic amplitudes in PCs upon cerebellar slice electrophysiology. These findings indicate a progressive synaptic reorganization in the degenerating cerebellum in SCA7. Whether this increase in inhibitory synapse function represents an attempt at compensation to alleviate excessive aberrant electrical activity or is driving the disease process in the cerebellum remains to be determined. However, there is reason to expect that increased inhibitory synapse formation resulting in excessive inhibitory synaptic input to PCs could be deleterious. In the *ax^J^* mouse, loss of function of ubiquitin-specific protease 14 results in increased levels of the GABA(A) receptor on PCs and correspondingly increased inhibitory postsynaptic currents (IPSCs), which may contribute to cerebellar dysfunction and degeneration^73^. Similarly, GABA transporter subtype 1 knock-out mice develop ataxia due to impaired uptake of GABA in the synaptic cleft, resulting in slower decay of IPSCs at both PC and cerebellar granule neuron synapses^74^. Although these findings suggest that increased inhibitory synapse numbers and inhibitory interneuron synaptic input in SCA7 PCs is part of the disease process, it will be necessary to modulate GABA(A) receptor activity in SCA7 mice using genetic and pharmacological interventions to determine if this alteration is compensatory or contributory to disease. Differentiating between these two scenarios will dictate whether an effective treatment strategy for SCA7 cerebellar degeneration will be to block or to enhance inhibitory synaptic transmission, and will have implications for therapy development for related polyQ SCAs.

## MATERIALS & METHODS

### Animals

SCA7-266Q mice (B6.129S7-Atxn7^tm1Hzo^/J; JAX#:008682) were maintained on a C57BL/6J background at Duke University DLAR facility and at UC Irvine ULAR facility. SCA3-Q84 mice (B6;CBA-Tg(ATXN3*)84.2Cce/IbezJ; JAX#:012705) were maintained on a C57BL/6J background at the University of Michigan. Mice were housed in groups of 2-5 per cage with a 12 hour dark-light cycle and free access to food and water. All animal work adhered to NIH guidelines under IACUC approved protocols at Duke, UC Irvine, University of Minnesota, University of Michigan, and University of Utah.

### snRNA-seq Tissue Preparation and Sequencing

Snap frozen cerebellar tissues from two male and two female SCA7-266Q mice and paired WT littermates were used for snRNA-seq experiments. Nuclear isolation with Purkinje nuclei enrichment was performed using a modified protocol based closely on Kozareva et.al. 2021^25^, by increasing K_2_SO_4_ concentration to 2.8g/500mL Dissociation Buffer which causes Granule Neuron nuclei to aggregate and fall out of solution. The resulting supernatant then contains a nuclear homogenate with elevated Purkinje cell proportions. Decreasing K_2_SO_4_ concentration to 1.4g/500mL in the Dissociation Buffer eliminated this effect. However, this concentration should be determined by the user, as other labs have observed varying K_2_SO_4_ levels to induce aggregates. After nuclei were purified, enriched for PCs, and stained with Hoechst 33342, separate wells representing nuclei from individual animals were separately labeled with MULTI-seq Cholesterol-Modified Oligos (CMOs) containing unique sample barcodes as described in McGinnis et.al. 2019^35^. Barcoded nuclei from each animal were then sorted into a single collection tube using a BD FACSAria II cell sorter from the Hoechst singlet population before loading onto the 10X Genomics Chromium Controller with the 3’ Gene Expression v3.1 kit. Sequencing libraries for snRNA and MULTI-seq barcodes were prepared separately from the cDNA as described in McGinnis et.al. 2019^35^, and sequenced on an Illumina Novaseq 6000. Identical processing steps were performed on post-mortem human cerebellar tissues (SCA7 n=5, Unaffected Controls n=4), but were instead sequenced on an Illumina Nextseq 2000.

### snRNA-seq QC and Analysis

Sequencing FASTQ files were demultiplexed before running through the 10X Genomics CellRanger *count* pipeline (v4.0.0) to generate a feature filtered barcode matrix containing UMI filtered gene counts for each cell barcode. The Seurat R package (v4.2.0) was used to normalize counts, identify variable features, scale data, and cluster nuclei with UMAP dimensional reduction. MULTI-seq barcode FASTQs were run through the deMULTIplex R package to generate sample specific barcode counts for each nucleus, remove doublets and ambiguous nuclei, and classify each nucleus to it’s animal of origin. Filtered and labeled nuclear clusters were then annotated to their cell type of origin using top cluster specific marker genes with Seurat::FindMarkers, referencing known cell type marker genes and the Mouse Transcriptomic Cerebellar Atlas^25^. The same processing steps were performed on post-mortem human cerebellar snRNA-seq data.

### Cell-type sub-clustering and Pseudobulk DEseq2 Differential Expression

Each annotated cell-type was further investigated by subsetting into individual Seurat objects before re-normalization, re-scaling, and re-clustering with UMAP reduction. Raw counts for each biological sample were pulled from the subset single-cell objects in separate columns of the count table for DEseq2, and differential expression was computed between groups of SCA7 and Control samples, regressing for sex effects in the model. DEGs were classified as significant if they met cutoff values of (p-adjusted < 0.05, Log2FoldChange > 0.5) for mouse samples and cutoff values of (p-adjusted < 0.05, Log2FoldChange > 2) for human samples. Cell Type DEG lists are provided in **Supplemental Data Table 1 (Mouse 5wk DEGs), Supplemental Data Table 2 (Mouse 8.5wk DEGs), and Supplemental Data Table 4 (Human DEGs)**.

For Human snRNA-seq, the number of cells recovered for each sample was much more variable than in mouse samples, so we focused our analysis to Granule Neurons, MLI-1, and MLI-2 clusters where cell recovery was highest. Samples contributing fewer than 25 cells to these cell types were excluded as replicates for differential expression analysis and count tables were normalized with a custom scaling factor (sample-cell-#/mean-cell-#) to account for gross differences in pseudobulk counts per sample.

### Zebrin-axis Quantification and Overlap with SCA7 PC DEGs

In Purkinje Cells we determined that there was significant divergence in SCA7 transcriptional profiles in populations of Zebrin-positive and Zebrin-negative PCs. To accurately characterize these cell populations, 8.5wk Wild-Type PC nuclei were subset into a new Seurat object before re-normalization, re-scaling, and re-clustering with UMAP reduction. UMAP clusters definitively characterized nuclei into clusters of ZebrinII+ or ZebrinII-PCs, which were compared using Seurat::FindMarkers (significance cutoffs: adjusted p_val < 0.05; |log2FC| > 0.25) to generate a gene list representing the Zebrin transcriptional axis (**Supplemental Data Table 3**). These Zebrin-axis genes were intersected with the SCA7 Purkinje Cell DEGs and statistical enrichment was calculated using a Hypergeometric test with a Bonferonni correction parameter of 13,078 matching the number of Gene Ontology Biological Process terms for multiple testing.

### Zebrin-axis WT and SCA7 Cross-comparison

The same steps described above to compute the WT Zebrin-axis were performed identically for 8.5wk SCA7 PC nuclei without any log2FC cutoffs. Differential expression outputs from WT and SCA7 Zebrin-axes were then directly compared by log2FC values for each gene and plotted together before running a linear regression to determine expression trends between WT and SCA7 Zebrin-axes.

### Quantitative Synapse IHC

To quantify cerebellar synapses, SCA7-266Q mice and WT littermates were perfused with 4% PFA after clearing with PBS and brains were post-fixed in 4% PFA overnight. Brains were then cryoprotected by soaking in 10% sucrose/PBS, 20% sucrose/PBS, and 30% sucrose/PBS in 24hr increments before mounting cerebellar hemispheres in OCT and snap-freezing in a liquid nitrogen isopentane bath. A Leica CM3050S Cryostat was used to prepare 30μm sagittal slices from the cerebellar vermis which were cryopreserved in 50% Glycerol: 30% Sucrose TBS. Synapse staining was then performed by washing slices 3X in TBST (0.2% Triton-X) for 10 minutes, blocking with 10% NGS in TBST at room temp for 1hr, and staining with primary antibody cocktail in 10% NGS-TBST overnight at 4°C. Slices were then washed 3X in TBST for 10 minutes, and stained with secondary fluorescent antibodies at a concentration of 1:300 for 2hrs in 10% NGS-TBST at room temp. Finally, slices were stained with Hoescht 33342 (1:5000) for 10 minutes in TBST and washed 2X for 10 minutes in TBST before mounting to microscope slides in Prolong Glass Antifade Mountant. Slides were imaged on a Zeiss 880 Inverted Confocal microscope at appropriate magnification (20X VGlut2 climbing fibers, 63X VGat-Gepherin inhibitory synapses) using Z-stack settings to collect 15 images with 0.34 μm step size in cerebellar Lobule VII. The PunctaAnalyzer ImageJ tool^75^ was used to separate the Z-stack images into 5 sets of 3 consecutive slices (∼1 μm total) and compute puncta overlap between pre-synaptic and post-synaptic markers, and averaging the co-localized synapse numbers from each slice to generate a final synapse count for the Field of View.

### Cerebellar Slice Electrophysiology

Purkinje Cell electrophysiology was performed in 250μm acute cerebellar slices in the coronal plane of the posterior vermis in Lobule VII as described in Tsai et.al. 2012^76^. Briefly, slices were cut in ice-cold Cutting Solution consisting of (mM): 130 K-Gluconate, 15 KCl, 0.05 EGTA, 20 HEPES, 25 Glucose (pH 7.4, osmolarity 320) before transferring to artificial cerebrospinal fluid (aCSF) containing (mM): 125 NaCl, 26 NaHCO_3_, 1.25 NaH_2_PO_4_, 2.5 KCl, 25 Glucose, 1 MgCl, 2 CaCl_2_ (pH 7.3, osmolarity 310). Slices were equilibrated with carbogen (95% O_2_ and 5% CO_2_) in aCSF for 20 minutes before moving to room temp with constant supply of carbogen. Recordings were taken in aCSF without addition of synapse or action potential inhibiting agents with borosilicate patch pipettes pulled to a resistance of 3-7 MΩ filled with an internal solution containing (mM): 140 Cs-Gluconate, 15 HEPES, 0.5 EGTA, 2 TEA-Cl, 2 MgATP, 0.3 NaGTP, 10 phosphocreatine-Tris_2_, 2 QX 314-Cl (pH 7.2). Action Potentials were recorded in a cell-attached configuration with a Gigaohm resistance seal before breaking into the membrane for whole-cell voltage clamp recordings of Spontaneous Post-Synaptic Currents. sEPSCs were recorded at -75mV and sIPSCs were recorded at +10mV. Membrane potentials were not corrected for the liquid junction potential.

### Zebrin-II IHC and Amplitude Analysis

Zebrin-II immunohistochemistry experiments were performed similarly as described above for Quantitative Synapse IHC with some modifications. Coronal 100μm slices were taken from the posterior cerebellum, and secondary antibody incubation time was increased to 4hrs. Images were taken at a 10X magnification on the Zeiss 880 Inverted Confocal microscope with a tile-scan to capture Zebrin-II and Calbindin immunofluorescence across the entire cerebellar slice. Zebrin striping amplitudes were then analyzed by taking xy-intensity values across the Purkinje and Molecular layer of the medial 3 Zb+ stripes (P1+,P2+) and medial 4 Zb-stripes (P1-,P2-) of cerebellar Lobule VIII. The signal was collapsed by taking the median intensity value along the x-axis, and a Loess polynomial regression was calculated to generate a signal trace across the region. Local minima and maxima were identified on the regression trace, and signal amplitudes were calculated for each consecutive stripe before computing the average Zebrin amplitude from these values.

### Quantitative Real-Time PCR (qRT-PCR)

Bulk RNA was purified from post-mortem human cerebellar tissues by pulverizing tissue with a cold mortar and pestle on dry ice. Roughly 25mg of pulverized tissue was then lysed with TRIzol (ThermoFisher) and purified with the Direct-zol RNA MiniPrep Plus kit (Zymo Research). Total RNA was then treated with TURBO DNase (ThermoFisher) and reverse transcribed to cDNA with SuperScript IV VILO (ThermoFisher). Taqman probes (ThermoFisher) against target genes with FAM dye were used to amplify cDNA in Taqman Gene Expression Master Mix (ThermoFisher) and compared to endogenous 18S rRNA with a Taqman VIC probe to generate Delta CT (dCt) values. Relative expression fold changes were calculated using the 2^-ddCt^ method against the mean dCt value for unaffected controls as reference. Two-tailed t-test was used to compute significance between SCA7 and unaffected controls.

## ACKNOWLEDGEMENTS

Anti-Zebrin-II antibody was generated and generously provided by Dr. Richard Hawkes and Dr. Carol Armstrong, University of Calgary. Post-mortem Human Tissues were provided by L. Ranum (Center for NeuroGenetics, University of Florida) and M. Perkins (University of Michigan Brain Bank). This work was supported by grants from the N.I.H. (Genomic Innovator Award to C.B.L.; R01 EY014061 and R35 NS122140 to A.R.L.S., R35 NS127253 and R37 NS033123 to SMP, R01 NS097903 to DRS, and R35 NS127248 to H.T.O), the Holland-Trice Scholars Award (L.C.B), and by the National Science Centre, Poland (NCN), project no UMO-2021/43/D/NZ3/03006 (P.M.S).

## Author Contributions

L.C.B. and A.R.L.S. provided the conceptual framework for the study. L.C.B, A.R.L.S, C.B.L, P.M.S., H.S.M., D.R.S., H.T.O., S.M.P., and C.H. designed the experiments. L.C.B., P.M.S., G.A., J.C., L.A.D., S.I.J., H.S.M., and D.R.S. performed the experiments. L.C.B., P.M.S., G.A., C.B.L., and A.R.L.S. analyzed the data. L.C.B, C.B.L, and A.R.L.S. wrote the manuscript.

## Declarations

The authors have nothing to declare.

## Supplementary Figure legends

**Supplemental Figure S1:**
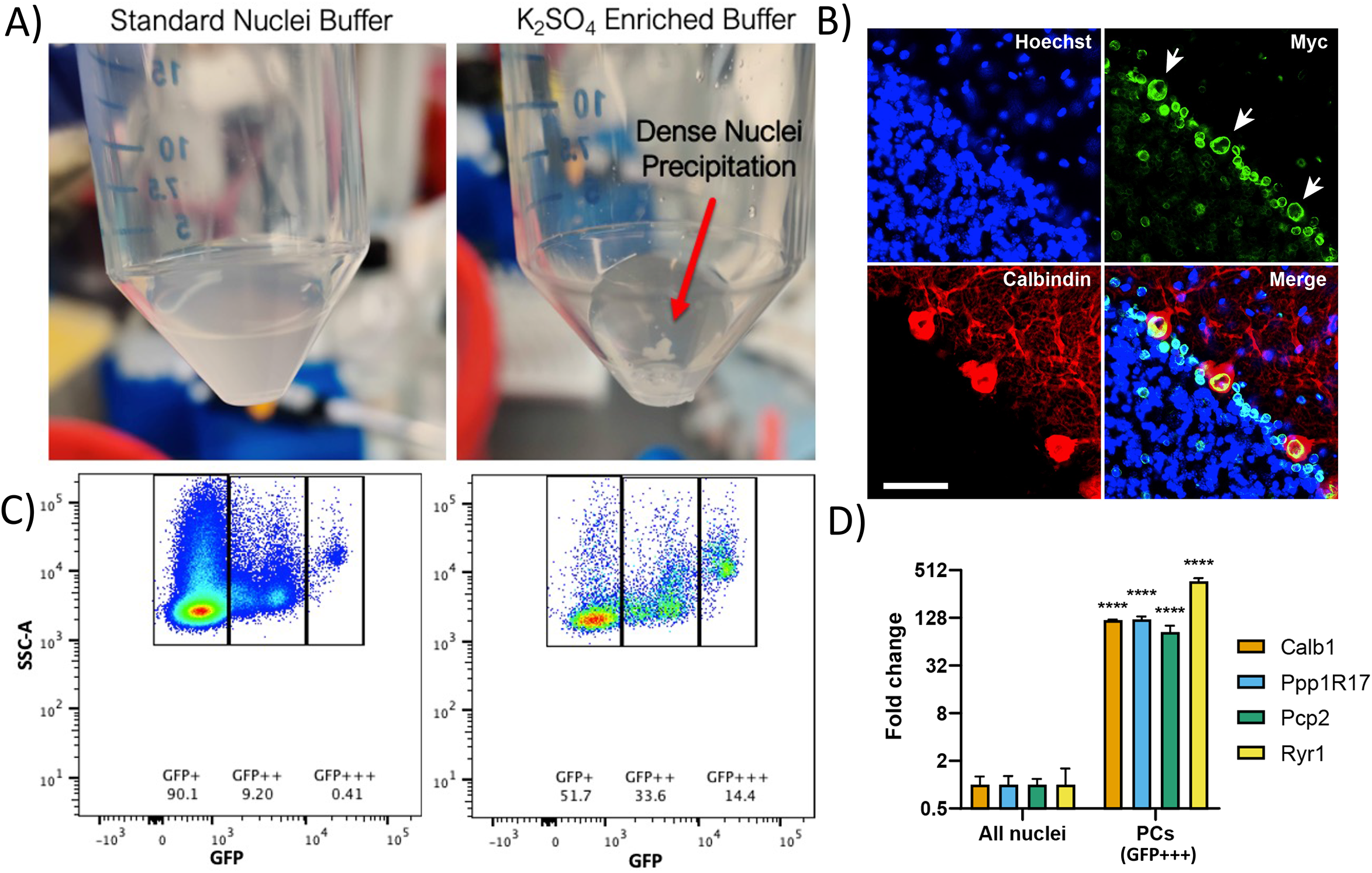
Purkinje Cell Nuclear Enrichment Method. A) Images of nuclear extraction buffers with standard concentration of potassium sulfate generating a homogenous suspension of cerebellar nuclei (left), and enriched concentrations of potassium sulfate leading to precipitation of granule neuron nuclei (right). B) Immunohistochemistry from sagittal cerebellar slices of INTACT Sun1-GFP mice display nuclear envelopes labeled with GFP. White arrows indicate Purkinje Cell nuclei which are notably larger than surrounding cell types. C) Flow Cytometry of nuclei isolated from INTACT Sun1-GFP mice with Standard and K_2_SO_4_ Enriched extraction buffers. Side-scatter (SSC-A) and GFP fluorescence separate cerebellar cell types by nuclear size and granularity. K_2_SO_4_-enriched extraction buffers result in a selective enrichment of GFP+++ nuclei. D) qRT-PCR from sorted total nuclei vs. GFP+++ gate validates enriched expression of Purkinje Cell markers (n=3).

**Supplemental Figure S2:**
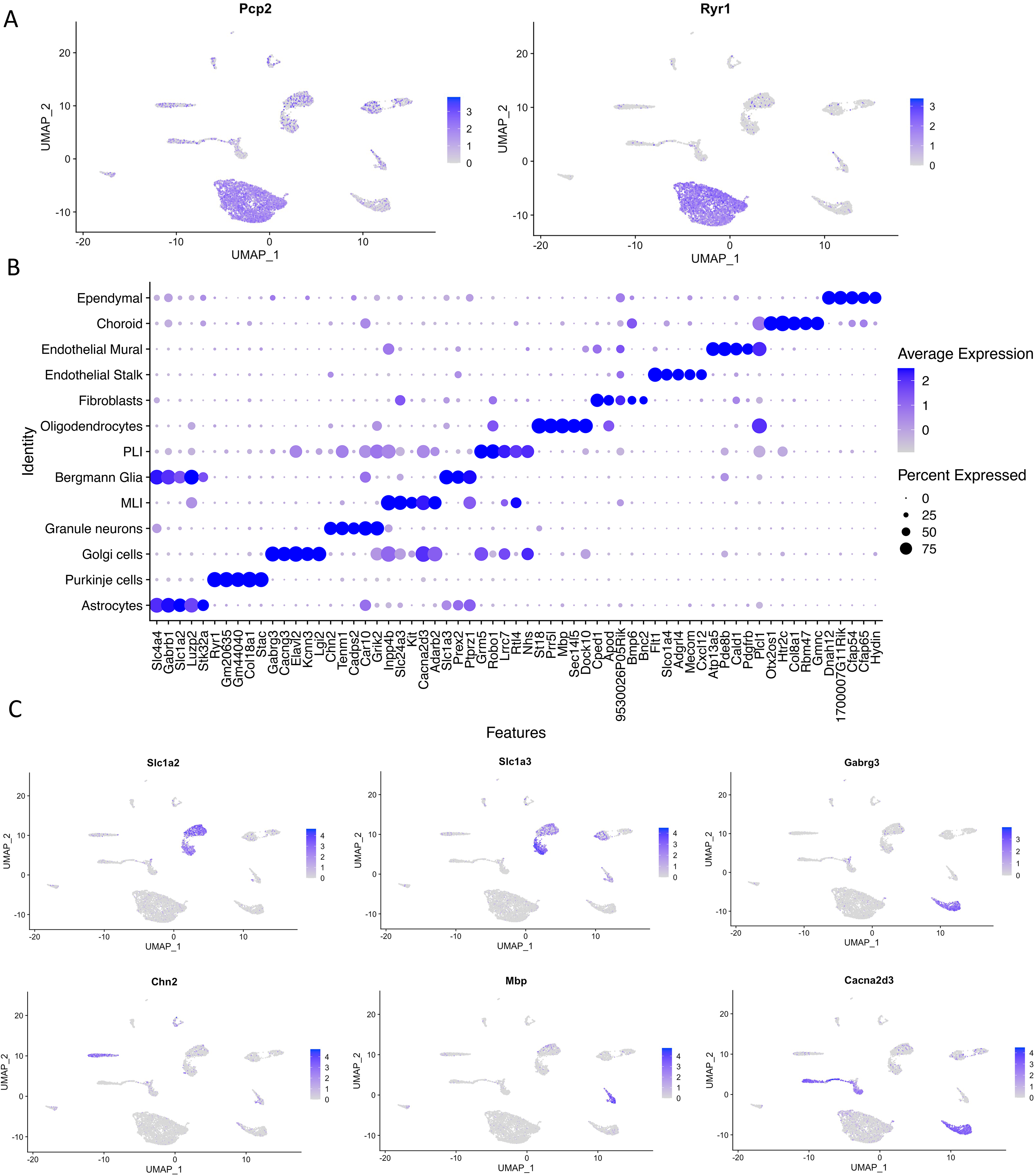
snRNA-seq Cell Type Marker Genes. A) Seurat::FeaturePlots showing canonical Purkinje Cell marker *Pcp2* and non-canonical marker *Ryr1* which showed highest specificity to Purkinje Cells in our dataset. B) Dot Plot showing top 5 marker genes for all cell types recovered. C) Feature plots visualizing cluster specific expression of top markers for Astrocytes (*Slc1a2*), Bergmann Glia (*Slc1a3*), Golgi Cells (*Gabrg3*), Granule Neurons (*Chn2*), Oligodendrocytes (*Mbp*), Molecular Layer Interneurons (*Cacna2d3*)

**Supplemental Figure S3:**
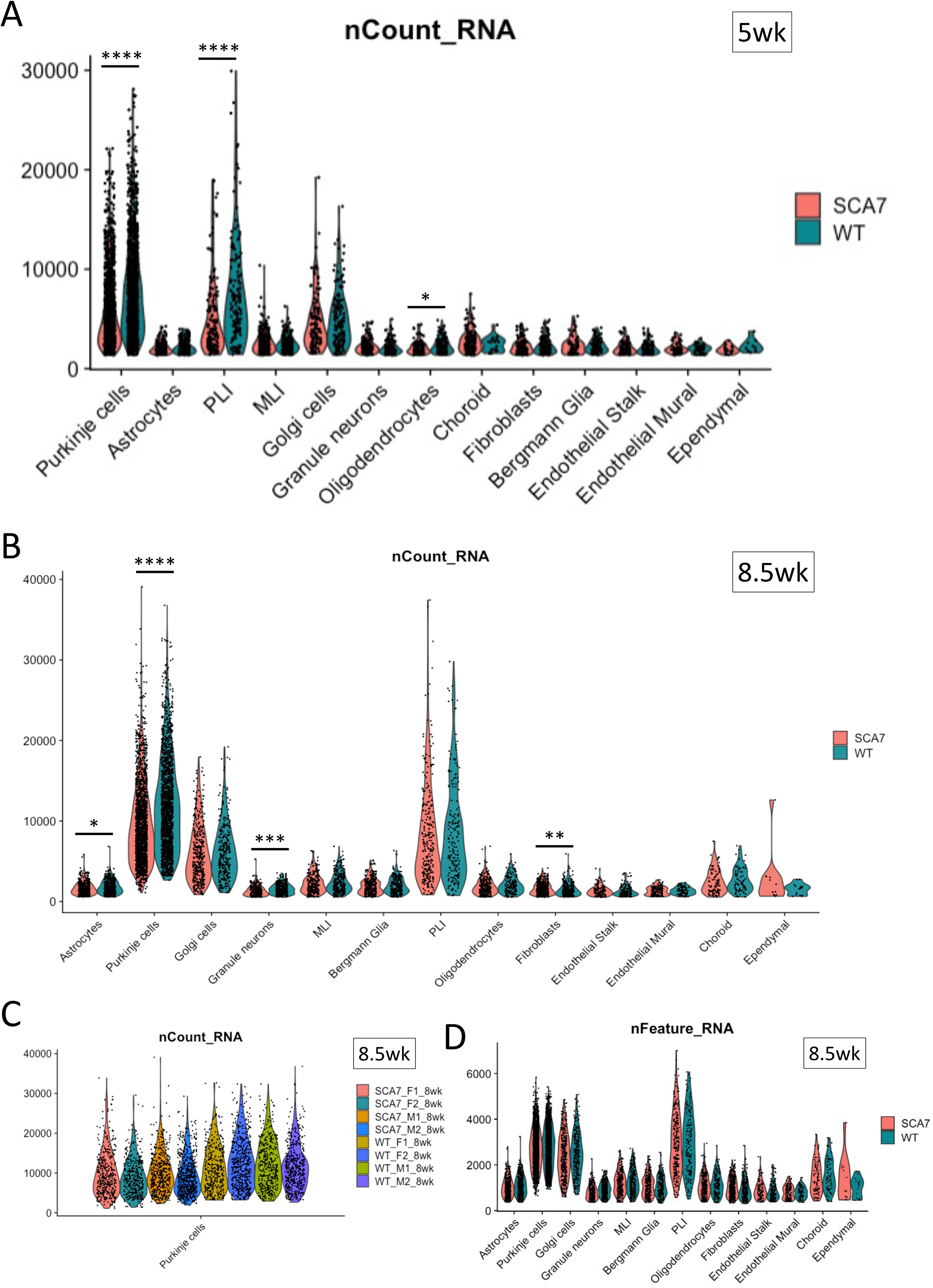
Cell Type Selective Decrease in RNA Abundance in SCA7-266Q snRNA-seq. A) Violin Plot showing RNA counts per nucleus (nCount_RNA) for the 5 week timepoint split by cell type and genotype. Statistical differences between SCA7 and WT were calculated for each cell type by Wilcoxon Rank Sum test. p-value cutoffs: * < 0.05, ** < 0.01, *** < 0.001, **** < 0.0001. B) Violin Plot showing RNA counts for the 8.5 week timepoint. C) Violin Plot showing RNA counts for 8.5 week Purkinje Cells split by animal. D) Violin Plot showing no significant difference in number of genes detected per nucleus (nFeature_RNA) for the 8.5 week timepoint split by cell type and genotype.

**Supplemental Figure S4:**
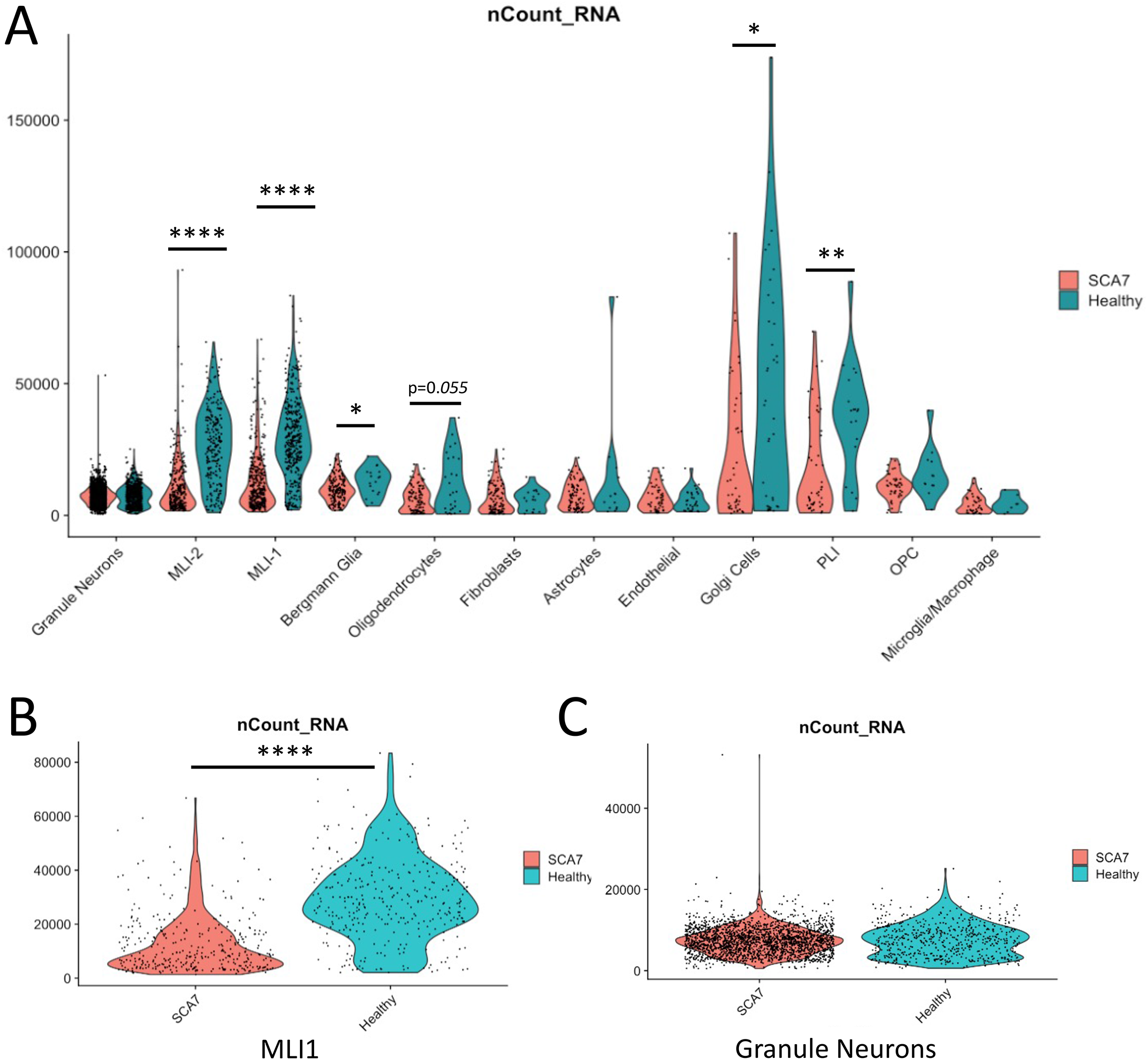
Cell Type Selective Decrease in RNA Abundance in Human SCA7 snRNA-seq. A) Violin Plot showing RNA counts per nucleus (nCount_RNA) for human Post-mortem Cerebellum snRNA-seq split by cell type and disease status. Statistical differences between SCA7 patients and unaffected controls were calculated for each cell type by Wilcoxon Rank Sum test. p-value cutoffs: *p < 0.05, **p < 0.01, ***p < 0.001, ****p < 0.0001. B) Violin Plot showing RNA counts for Type-I Molecular Layer Interneurons grouped by disease status. C) Violin Plot showing RNA counts for Granule Neurons grouped by disease status.

**Supplemental Figure S5:**
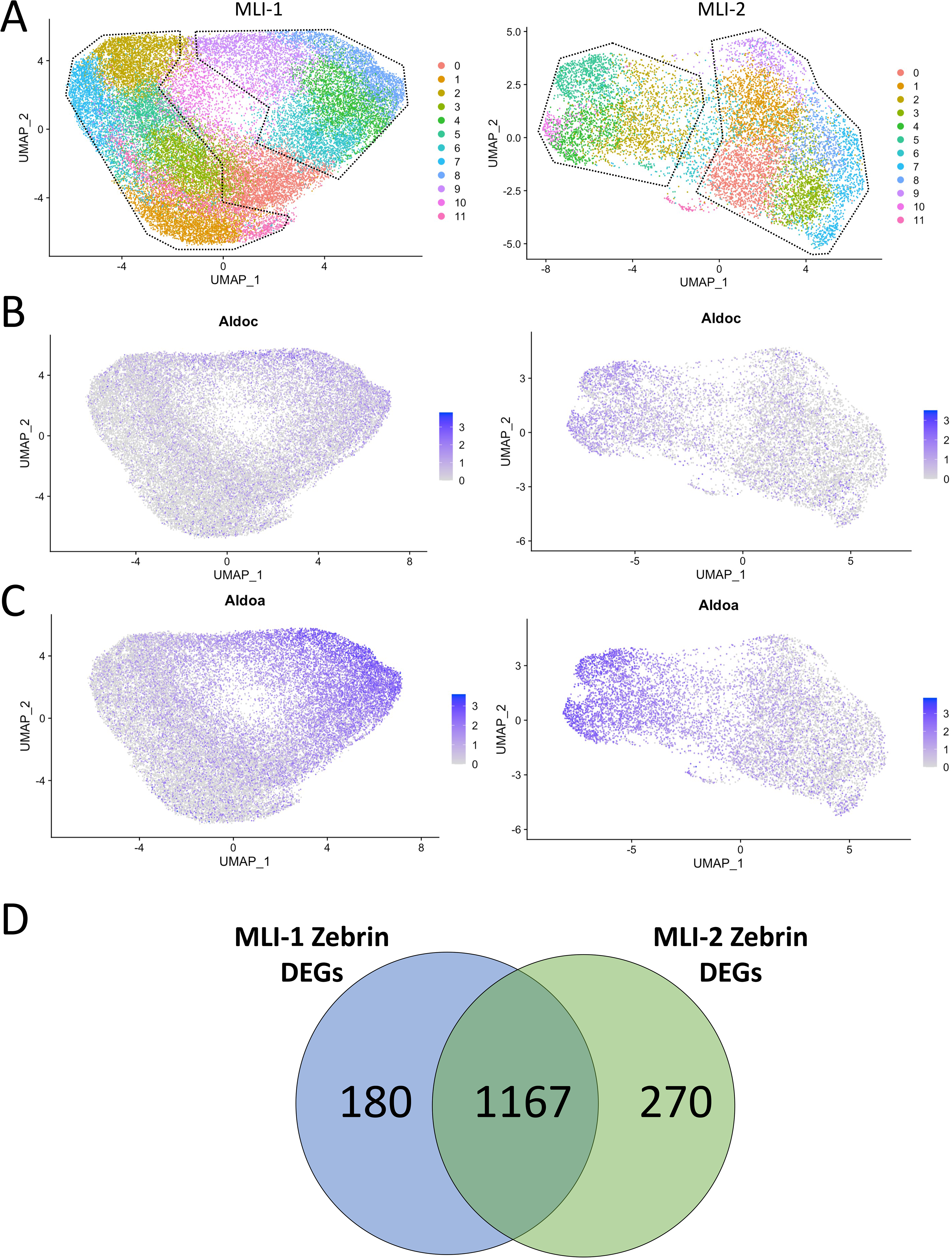
Subclusters of MLI-1 and MLI-2 Nuclei from Transcriptomic Mouse Cerebellar Atlas Show Zebrin-II Gene Expression Patterning. A) UMAP plots showing re-clustered MLI-1 and MLI-2 nuclei from Kozareva et al. transcriptomic Mouse Cerebellar Atlas. Dotted lines highlight *Aldoc/Aldoa-*‘high’ and *Aldoc/Aldoa-*‘low’ subclusters. B) Feature Plots showing normalized expression of *Aldoc* in MLI-1 and MLI-2 clusters. C) Feature Plots showing normalized expression of *Aldoa* in MLI-1 and MLI-2 clusters. D) Venn-Diagram showing overlap of Zebrin Gradient DEGs between *Aldoc/Aldoa-*‘high’ and *Aldoc/Aldoa-*‘low’ subclusters in MLI-1 and MLI-2.

**Supplemental Figure S6:**
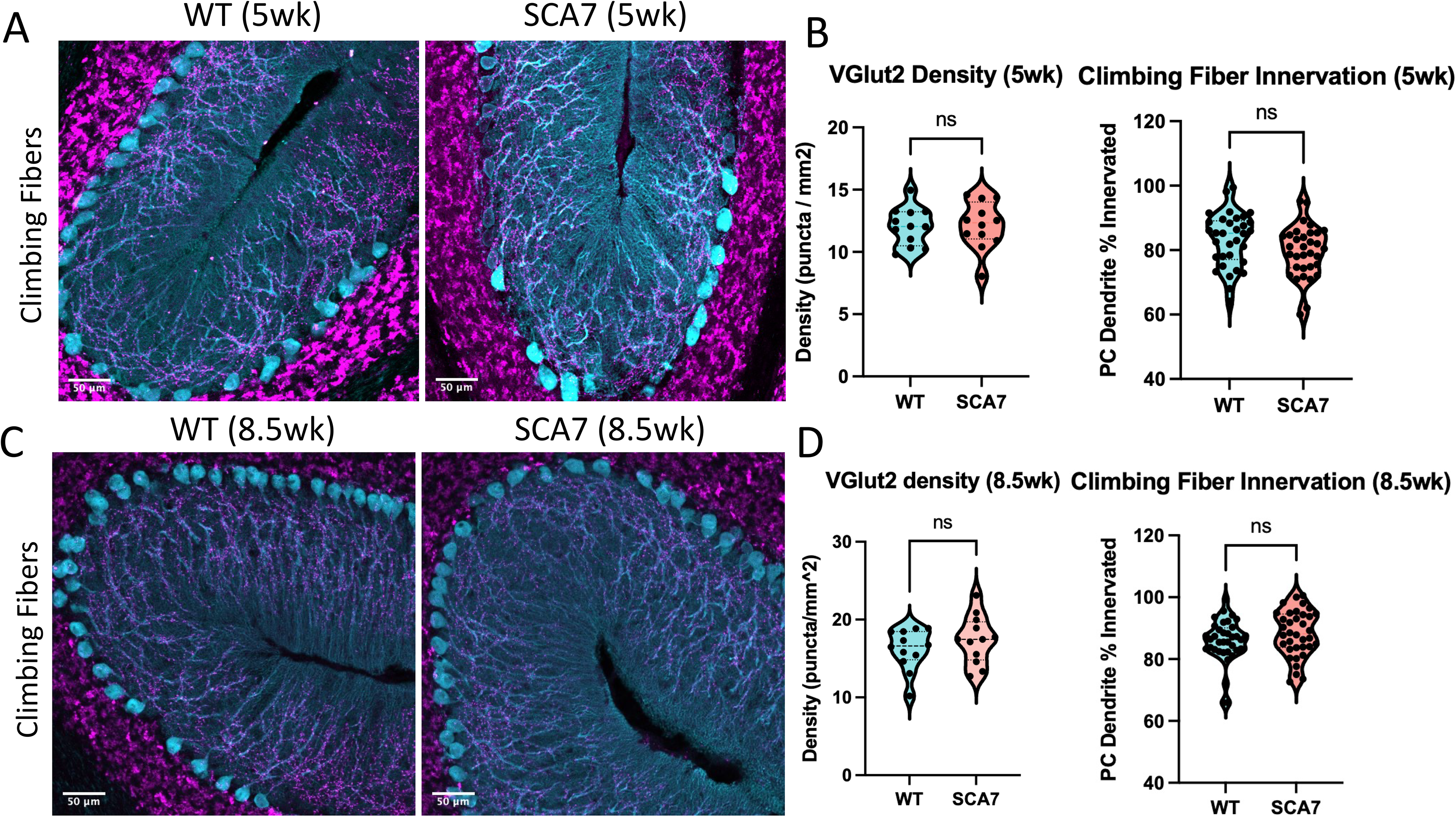
VGlut2 Climbing Fiber Synapses Remain Unchanged in Symptomatic SCA7-266Q Animals. A) Quantitative Synapse immunohistochemistry representative images for excitatory climbing fiber synapses in cerebellar Lobule VII in 5 week-old mice. 20X magnification; magenta = VGlut2; cyan = Calbindin-1. B) Quantification of 5 week climbing fiber synapses by puncta density and percent innervation along full length of the PC dendrite. n=6 mice/genotype; two images per animal. Statistics calculated with two-tailed t-test, p=n.s. C) Quantitative Synapse immunhistochemistry representative images for excitatory climbing fiber synapses in cerebellar Lobule VII in 8.5 week-old mice. 20X magnification; magenta = VGlut2; cyan = Calbindin-1. D) Quantification of 8.5 week climbing fiber synapses by puncta density and percent innervation along full length of the PC dendrite. n=6 mice/genotype; two images per animal. Statistics calculated with two-tailed t-test, p=n.s.

